# Glycogen metabolism acts in neurons to support glycolytic plasticity

**DOI:** 10.1101/2025.04.10.648039

**Authors:** Milind Singh, Aaron D. Wolfe, Anjali A. Vishwanath, Anastasia Tsives, Ian J. Gonzalez, Sarah E. Emerson, Richard H. Goodman, Daniel A. Colón-Ramos

**Affiliations:** Department of Neuroscience and Department of Cell Biology, Yale University School of Medicine; New Haven, CT 06536, USA; Wu Tsai Institute, Yale University; New Haven, CT 06510, USA

## Abstract

Glycogen is the largest energy reserve in the brain, but the specific role of glycogen in supporting neuronal energy metabolism *in vivo* is not well understood. We established a system in *C. elegans* to dynamically probe glycolytic states in single cells of living animals via the use of the glycolytic sensor HYlight and determined that neurons can dynamically regulate glycolysis in response to activity or transient hypoxia. We performed an RNAi screen and identified that PYGL-1, an ortholog of the human glycogen phosphorylase, is required in neurons for glycolytic plasticity. We determined that neurons employ at least two mechanisms of glycolytic plasticity: glycogen-dependent glycolytic plasticity (GDGP) and glycogen-independent glycolytic plasticity (GIGP). We uncover that GDGP is employed under conditions of mitochondrial dysfunction, such as transient hypoxia or in mutants for mitochondrial function. We find that the ability of neurons to plastically regulate glycolysis through cell-autonomous GDGP is important for sustaining the synaptic vesicle cycle. Together, our study reveals that, *in vivo*, neurons can directly use glycogen as a fuel source to sustain glycolytic plasticity and synaptic function.

## Introduction

Energy consumption in the brain is tightly linked to its activity states (Attwell & Laughlin, 2001; Dienel, 2019; Vaishnavi et al., 2010). The link between energy consumption and brain activity states is arguably best understood in the context of functional brain imaging studies, in which glucose consumption is used as a proxy for underlying changes in local neuronal activity (Bingham et al., 2005; Macapinlac, 2006). From these studies, we know that energy metabolism is plastically modulated in specific regions of the brain and in response to spatiotemporal changes in brain activity states (Hyder & Rothman, 2017; Vergara et al., 2019). Conditions that alter the metabolic state of the brain, such as hypoxia, starvation and hypoglycemia, have profound effects on cognitive functions, further underscoring the tight relationship between the regulation of energy metabolism and brain function (Cherubini et al., 1989; Languren et al., 2013; Li et al., 2018). Cell biological studies have also revealed that neurons dynamically regulate their energy metabolism to meet changes in activity states (Díaz-García et al., 2017; Wolfe et al., 2024). Therefore an emerging and supported concept in neuroscience, both from functional brain imaging and cell biological studies, is that energy metabolism dynamically adapts in specific brain tissues to support brain function (Bordone et al., 2019; Rae et al., 2024).

At a cell biological level, energy metabolism can be regulated primarily via the modulation of mitochondrial function and glycolytic plasticity (Rangaraju et al., 2014; Vergara et al., 2019; Watts et al., 2018). While mitochondrial respiration and glycolysis are well-conserved pathways present in most cells, the specific regulation of these energy-producing pathways varies across cell types and brain regions (Magistretti & Allaman, 2015). The spatiotemporal modulation of these energy-producing pathways endows the brain with the capacity to sustain function during varying environmental conditions, including alterations in dietary nutrients or transient hypoxia (Camandola & Mattson, 2017; Shetty et al., 2012). Because neuronal computations are energetically expensive and rapidly change across specific brain regions, metabolic plasticity allows the brain to adjust energy metabolism to meet changes in activity states (Attwell & Laughlin, 2001; Vergara et al., 2019). Despite their importance in sustaining brain function, the underlying regulation of metabolic plasticity of the brain, particularly in neurons, is not well understood.

The relative abundance of different fuels for energy production influences the contribution of specific and distinct energy-producing pathways to cells (Yellen, 2018). This is arguably best understood in muscles, in which the differential use of glucose, glycogen, fat stores, or proteins as fuels preferentially drives glycolysis or mitochondrial respiration (Blomstrand & Saltin, 1999; Hargreaves & Spriet, 2020). The preferential use of glycolysis or mitochondrial respiration then influences the physiological properties of the muscle fibers, including contractile speeds and fatigue (Hargreaves, 2000; Hargreaves & Spriet, 2020; Smith et al., 2023). While our understanding of fuel usage in neurons, and how it influences function in specific contexts, is less characterized, it has been proposed that neurons use lactate as a primary fuel source, with glia preferentially performing glycolysis to supply lactate for neuronal respiration (Magistretti & Allaman, 2015, 2018). Recent reports have also demonstrated that neurons perform glycolysis, regulate glucose uptake based on neuronal activity states, and modulate glycolytic plasticity to meet energy demands (Ashrafi et al., 2017; Díaz-García et al., 2017; Wolfe et al., 2024). The fuel sources used by neurons in specific contexts, and how these fuel sources influence neuronal functions and metabolic plasticity are active areas of research.

Glycogen is the largest energy reserve in the brain (Bélanger et al., 2011). Glycogen is a multibranched polysaccharide composed of glucose molecules, and it is the main form of glucose storage in animals (Roach et al., 2012). The role of glycogen in energy metabolism is best understood for muscles, where glycogen granules act as ‘capacitors’, buffering changes in carbon flux and energy utilization inside cells (Prats et al., 2018). In the nervous system, glycogen plays neuroprotective roles, and disruptions in glycogen utilization in the brain have been associated with alterations in memory formation and susceptibility to epilepsy (Duran et al., 2013; Duran, Gruart, Varea, et al., 2019; Duran et al., 2020; Gibbs et al., 2006). While brain glycogen is predominantly stored in astrocytes, neurons also express the enzymes necessary for the formation and breakdown of glycogen (Duran, Gruart, López-Ramos, et al., 2019; Saez et al., 2014; Vilchez et al., 2007). It was recently demonstrated that neurons synthesize this polysaccharide and that the availability of glycogen in neurons is necessary for their tolerance to hypoxia (López-Ramos et al., 2015; Saez et al., 2014). Moreover, mice with a neuron-specific deficiency in producing glycogen displayed specific deficiencies in long-term potentiation (LTP) and learning, underscoring the neuron-specific requirements of glycogen (Duran, Gruart, Varea, et al., 2019). These findings support the role of glycogen biosynthetic pathways in neuronal function and underscore the importance of elucidating how glycogen contributes to metabolic plasticity in neurons.

In this study, we established a system in *C. elegans* to probe glycolytic plasticity in single cells of living animals via the use of the glycolytic biosensor HYlight (Koberstein et al., 2022). HYlight monitors levels of fructose 1,6 bisphosphate (FBP), a metabolite produced by the rate-limiting enzyme phosphofructokinase, which represents a committed step of glycolysis (Koberstein et al., 2022). By adapting microfluidic devices (Miller et al., 2005; Albrecht & Bargmann, 2011; Jang et al., 2021a) that allow us to induce energy stress *in vivo* while monitoring HYlight, we determine that cells dynamically adjust their glycolytic states to meet transient changes in energy demands. To identify mutants that affect the dynamic regulation of glycolytic plasticity, we performed an RNAi screen and determined that PYGL-1, an ortholog of glycogen phosphorylase, is cell-autonomously required for metabolic plasticity in neurons. We uncovered two mechanisms of glycolytic plasticity which are deployed in specific contexts: glycogen-dependent glycolytic plasticity (GDGP) and glycogen-independent glycolytic plasticity (GIGP). We find that GDGP is preferentially employed under conditions of mitochondrial dysfunction, such as transient hypoxia or in mutants for mitochondrial function, and that this ability of neurons to plastically regulate glycolysis through cell-autonomous GDGP is important for sustaining synaptic function. Together, our study supports that: 1) neurons can utilize glycogen as a cell-autonomous fuel source, and 2) glycogen sustains context-specific regulation of glycolytic plasticity and synaptic function.

## Results

### RNAi screen uncovers the role of Glycogen phosphorylase/PYGL-1 in regulating glycolytic plasticity

Cells dynamically regulate their glycolytic states to meet changing energy demands. To understand how cells plastically regulate glycolytic transients, we examined glycolysis *in vivo* by using HYlight, a biosensor for Fructose-1,6-bisphosphate (FBP) (Koberstein et al., 2022). FBP was chosen because it represents the committed step in glycolysis, produced through the actions of phosphofructokinase (PFK-1.1 in *C. elegans*). HYlight is a ratiometric sensor that dynamically measures FBP levels and enables *in vivo* probing of changes in the glycolytic states of cells (Koberstein et al., 2022; Wolfe et al., 2024). To alter energy stress, we used a custom-built microfluidic device that enabled precise control of oxygen levels while imaging live animals (Jang et al., 2021b). Because oxygen is necessary for oxidative phosphorylation in mitochondria, transient hypoxia inhibits mitochondrial metabolism to induce energy stress. Our microfluidic device enables minute-scale changes in transient hypoxic or normoxic conditions in living animals, enabling dynamic *in vivo* control of mitochondria function in ways that are not possible with other pharmacological or genetic approaches (Figure 1A) (Jang et al., 2021).

**Figure 1:**
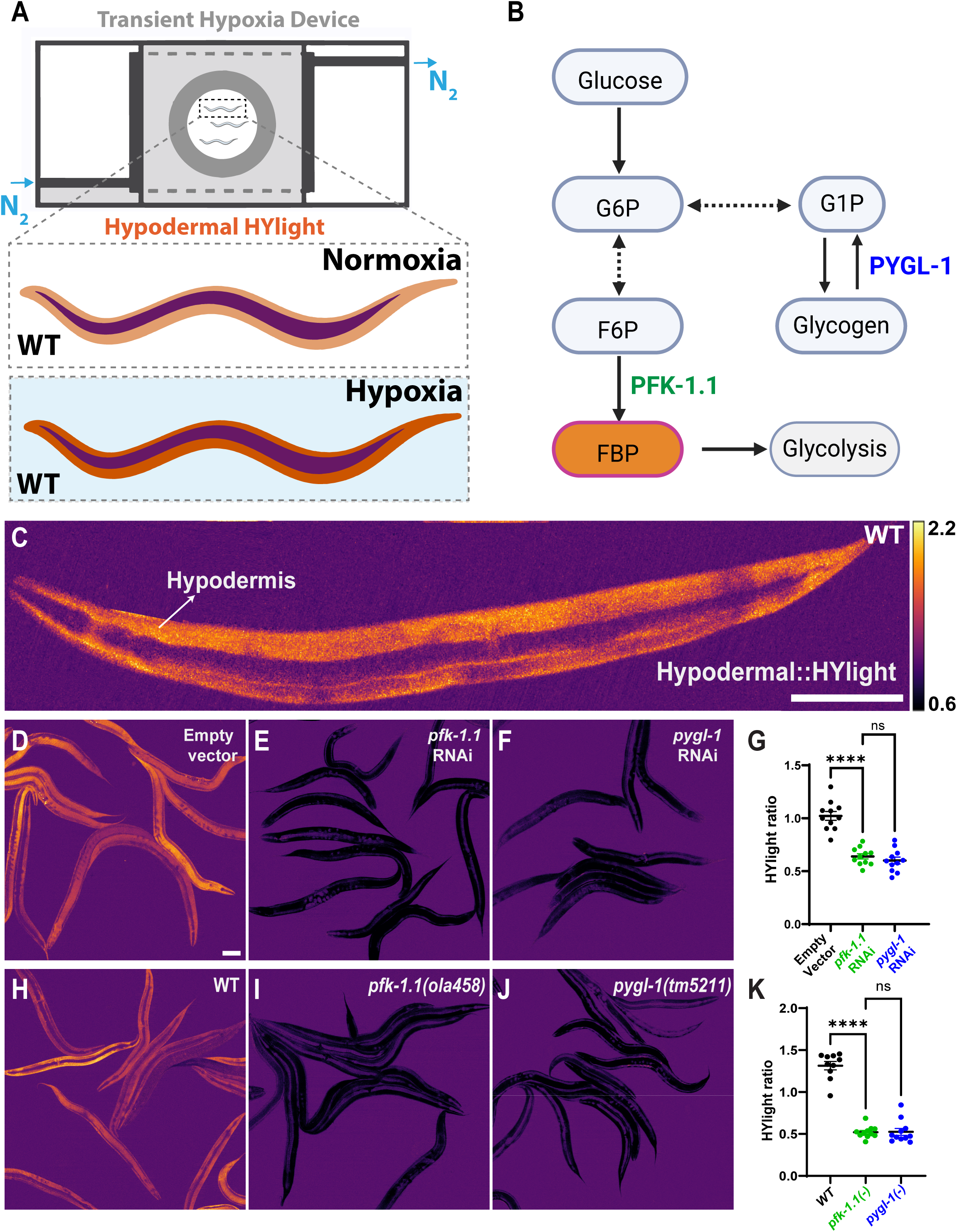
Identification of *pygl-1/*Glycogen phosphorylase as a regulator of glycolysis in the *C. elegans* hypodermis. **(A)** Schematic of the microfluidic device used in this study to induce transient hypoxia via controlled flowing of nitrogen gas (N_2_) (Albrecht & Bargmann, 2011; Jang et al., 2021a; Miller et al., 2005). *C. elegans* expressing the HYlight biosensor in the hypodermis were used in this figure. Under hypoxic conditions, oxidative phosphorylation is inhibited, glycolytic flux increases and the HYlight signal increases, as shown in schematic and previously described (Koberstein et al., 2022) . **(B)** Diagram depicting the connection between glycolysis and glycogenolysis. PYGL-1, the *C. elegans* ortholog of glycogen phosphorylase, breaks down glycogen to generate glucose-1-phosphate, which is converted into glucose-6-phosphate and enters the glycolytic pathway. For an explanation of the RNAi screen, please see Supplementary Figure 1-S2. **(C)** Ratiometric image of HYlight expressed in the hypodermis, showing the emission ratio after 488 nm and 405 nm excitation. The calibration bar indicates ratiometric signal intensity. **(D–G)** Hypodermal HYlight ratios in normoxia for WT worms fed empty vector (D), *pfk-1.1* RNAi (E) , or *pygl-1* RNAi (F) and quantification (G). In (G), each dot represents one worm; **** indicates *p* < 0.0001, calculated using Brown-Forsythe and Welch’s ANOVA followed by Dunnett’s multiple comparisons test. Eleven animals were analyzed per condition. **(H–K)** Ratiometric images of hypodermal HYlight in normoxia in WT (H), *pfk-1.1(ola458)* (I), and *pygl-1(tm5211)* (J) mutant animals and quantification (K). In (K), each dot represents one worm; **** indicates *p* < 0.0001, calculated using Brown-Forsythe and Welch’s ANOVA followed by Dunnett’s multiple comparisons test. Eleven animals were analyzed for *pfk-1.1(ola458)* and *pygl-1(tm5211)*, and ten for WT. Scale bar in C and D is 100µm. Scale bar in D corresponds to (D-J).

We first expressed the HYlight biosensor using a hypodermis-specific promoter (*col-19p*) (Figure 1-S1). We focused on the *C. elegans* hypodermal tissue because 1) the hypodermis is a central hub for the regulation of metabolism in *C. elegans* (Dowen et al., 2016), 2) it expresses regulators of metabolism (Fukuyama et al., 2015) and 3) might be sensitive to disruptions of metabolic states (Kaletsky et al., 2018). From a technical standpoint, the hypodermis is also readily amenable to screens using RNAi libraries (unlike neurons, which require specialized genetic backgrounds for RNAi (Firnhaber & Hammarlund, 2013; Tavernarakis et al., 2000; Timmons et al., 2001).

Expression of HYlight in hypodermal cells displayed high HYlight signal levels, consistent with the hypodermis being a metabolically active cell with high levels of glycolysis, even at resting states (Figure 1C). We had previously examined HYlight in neurons of mutants for Phosphofructokinase, the enzyme that makes FBP, and demonstrated that in *pfk-1.1(ola458)* null mutant animals, HYlight signal is abrogated (Wolfe et al., 2024). To determine if the observed HYlight signal in the hypoderm was due to cellular glycolytic states, and to validate the use of RNAi to screen for mutants that affect glycolytic plasticity, we performed RNAi against PFK-1.1 by feeding worms bacteria expressing *pfk-1.1* dsRNA (Figure 1D-G). We observed that the HYlight signal in hypodermal cells of animals fed with *pfk-1.1* dsRNA was lower than that of vector control, consistent with the observed HYlight signal corresponding to glycolytic states (Figure 1D-E and 1G). Moreover, *pfk-1.1* RNAi animals that were placed in a microfluidic device and exposed to transient hypoxia did not display increases in HYlight signal, indicating an inability to adjust FBP levels in the absence of *pfk-1.1*, as expected (Figure 1-S1, compare to *pfk-1.1* null allele in Figure 1-S3). Together, these observations indicated that the hypodermis is a glycolytically active tissue that requires PFK-1.1 for glycolytic plasticity in response to transient hypoxia. Importantly, these experiments demonstrate that we can make use of HYlight, microfluidic devices and RNAi screens to identify regulators of cellular glycolytic plasticity.

To identify genes necessary for regulating metabolic states in single cells, we performed a targeted RNAi screen on available selected genes involved in metabolite transport, carbohydrate metabolism and fat metabolism (Figure 1-S2). We were interested in identifying genes that abrogated the ability of the organism to dynamically adjust its cellular glycolytic states upon the energy stress during transient hypoxia. We observed that *pygl-1* RNAi reduced FBP levels, similar to *pfk-1.1* RNAi (Figure 1E-G) and was also unable to dynamically adjust its glycolytic state upon transient hypoxia (Figure 1-S3). PYGL-1 is an ortholog of human PYGM (glycogen phosphorylase), an enzyme that cleaves glycogen to release glucose-1-phosphate, which then enters the glycolytic pathway (Figure 1B) (Agius, 2015). According to CeNGEN single-cell transcriptomics data, *pygl-1* is ubiquitously expressed across all major tissues in the worm, including neurons (Hammarlund et al., 2018).

To further examine our RNAi screen findings, we visualized HYlight levels in a previously-characterized loss-of-function mutation of *pygl-1, pygl-1(tm5211)*(Chen et al., 2023). Similar to the *pygl-1* RNAi, HYlight ratios in *pygl-1(tm5211)* mutants were lower than wild-type (WT) and phenocopied FBP levels observed in a *pfk-1.1(ola458)* null mutants (Figure 1H-K). Importantly, unlike wild-type controls, *pygl-1(tm5211)* were incapable of adjusting glycolytic states in response to transient hypoxia (Figure 1-S3). Thus, PYGL-1, a glycogen breakdown enzyme, contributes to maintaining FBP levels in the worm hypodermis and is required for dynamic changes in glycolytic states in response to transient hypoxia.

### PYGL-1 is required in neurons to support glycolytic plasticity during hypoxia

Neurons can dynamically adjust glycolysis in response to energy demands such as transient hypoxia (Díaz-García et al., 2017; Wolfe et al., 2024), and this ability to adjust glycolytic metabolism to changing metabolic demands is termed glycolytic plasticity (Fendt et al., 2020; Papadaki & Magklara, 2022). To determine the role of glycogen in the regulation of glycolytic plasticity in neurons, we expressed the HYlight biosensor in *C. elegans* neurons using the *rab-3* panneuronal promoter, and measured FBP levels within the *C. elegans* nerve ring (Figure 2A). We observed that under resting conditions, neuronal metabolic states were cell-specific and diverse, consistent with previous reports (Figure 2B) (Wolfe et al., 2024). Upon transient hypoxia, we observed a concomitant increase in HYlight signal in the nerve ring, consistent with neurons dynamically adjusting their glycolytic state to meet the energy stress induced by transient hypoxia (Figure 2B’ and Movie S1). This increase was abrogated in *pfk-1.1(ola458)* null animals (Figure 2C-C’). We next measured panneuronal HYlight in the *pygl-1(tm5211)* mutant background and observed that under resting conditions, the HYlight ratios in the nerve ring were significantly lower in the *pygl-1(tm5211)* mutant as compared to WT animals but higher than in *pfk-1.1* mutants (Figure 2D and 2E). When *pygl-1(tm5211)* mutants were exposed to transient hypoxia, FBP levels did not increase, similar to what we observe in *pfk-1.1* mutants and in contrast to what we observed in WT worms (Figure 2D’, Figure 2F and Movie S2). This suggests that *pygl-1* is required for neurons to increase glycolytic flux in response to hypoxia.

**Figure 2:**
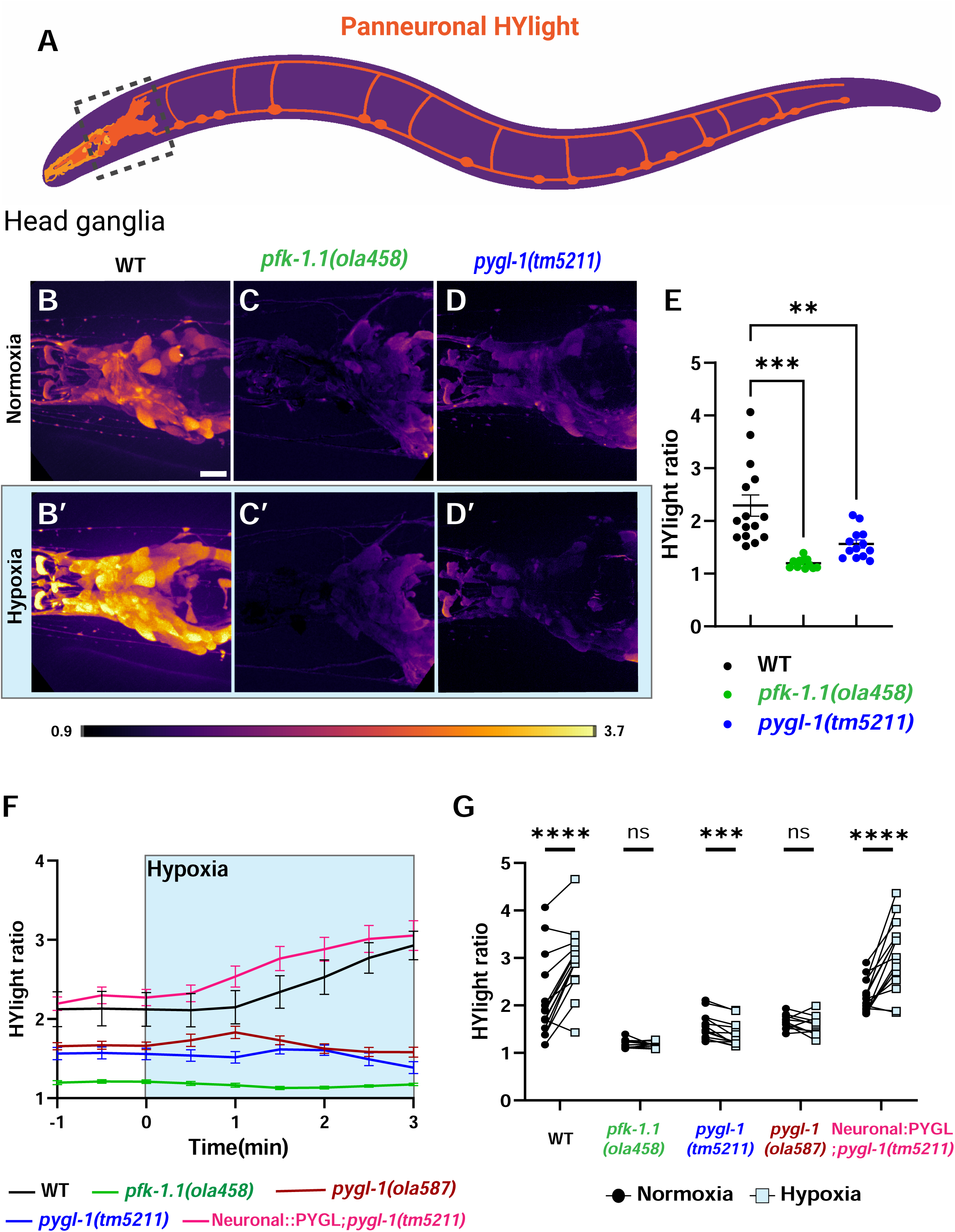
*pygl-1/*Glycogen phosphorylase sustains glycolysis in neurons during energy stress. **(A)** Schematic of a worm expressing HYlight in the nervous system using the panneuronal promoter *rab-3p*. Dashed box represents the images regions of the nerve ring (the head ganglia) imaged and quantified in the rest of the figure. **(B–D)** HYlight ratio (488/405 nm excitation) in the nerve ring of WT (B), *pfk-1.1(ola458)* null (C), and *pygl-1(tm5211)* mutants (D) during normoxia. **(B′–D′)** As (B-D), but after 3 minutes of transient hypoxia. **(E)** Quantification of HYlight ratios in the head ganglia under normoxia for WT, *pfk-1.1(ola458)*, and *pygl-1(tm5211)* animals. Each dot represents one worm; *** indicates *p* < 0.001, calculated using Brown-Forsythe and Welch’s ANOVA followed by Dunnett’s multiple comparisons test. Fifteen animals were analyzed for WT, 12 for *pfk-1.1(ola458)*, and 13 for *pygl-1(tm5211)*. **(F)** Plot of HYlight ratios over time in normoxia and hypoxia (hypoxia period shaded blue) for WT, *pfk-1.1(ola458)*, *pygl-1(ola587)*, *pygl-1(tm5211)*, and panneuronal rescue of PYGL-1A in *pygl-1(tm5211)* mutants. Error bars represent the standard error of the mean at each time point. Fifteen animals were analyzed for WT, 12 for *pfk-1.1(ola458)*, 13 for *pygl-1(tm5211)*, 11 for *pygl-1(ola587)*, and 15 for the neuronal PYGL-1A rescue in *pygl-1(tm5211)*. **(G)** Graph showing the change in HYlight ratios from normoxia to hypoxia for each genotype using data from (F). Normoxic values were taken at -1 minute, and hypoxic values at 3 minutes into hypoxia, as indicated in (F). *** denotes *p* < 0.001; **** denotes *p* < 0.0001, calculated using paired *t*-tests. Scale bar in B is 10µm. Scale bar in B corresponds to (B-D’).

The loss of glycolytic plasticity in neurons observed for *pygl-1* mutants could be due to a cell-autonomous role of glycogen phosphorylase in neurons, or a non-autonomous function of glycogen phosphorylase in other tissues. To establish the site of action of *pygl-1*, we conducted a tissue-specific rescue in the neurons of *pygl-1(tm5211)* mutants. We expressed the cDNA of the PYGL-1A isoform in neurons using the *rab-3* promoter in the background of *pygl-1(tm5211)*. This approach restored the disrupted FBP response phenotype during hypoxia to WT levels in the nerve ring (Figure 2F). These results support a cell-autonomous role for *pygl-1* in regulating glycolytic plasticity in neurons. To further validate the role of *pygl-1*, we generated a full gene deletion allele using CRISPR/Cas9, *pygl-1(ola587)*. Like the *pygl-1(tm5211)* allele, *pygl-1(ola587)* mutants failed to increase FBP levels during hypoxia (Figure 2F), confirming that *pygl-1* function is important for the neuronal glycolytic response to energy stress.

We quantified the change in FBP levels from normoxia to hypoxia across genotypes by calculating the difference in HYlight ratios before and after hypoxia. WT animals exhibited a significant increase in HYlight ratios, whereas *pfk-1.1(ola458)*, *pygl-1(tm5211)*, and *pygl-1(ola587)* mutants showed no increase. This defect was rescued by neuronal expression of PYGL-1A (Figure 2F and 2G). These results indicate that PYGL-1 is acting in the nervous system to support glycolytic plasticity during transient hypoxia. We term this requirement “glycogen-dependent glycolytic plasticity” (GDGP). Our results are consistent with studies that show that neurons have an active glycogen metabolism that protects cultured neurons from hypoxia-induced death and Drosophila from hypoxia-induced stupor(Saez et al., 2014). We now extend these findings, showing that glycogen breakdown is required for glycolytic plasticity in neurons during transient hypoxia *in vivo*, and suggesting that neurons can utilize glycogen as a fuel source during energetic stress.

### PYGL-1 is required in distinct neuron types to support glycolytic plasticity during hypoxia

Neuronal metabolic states are cell-specific and diverse across the nervous system (Bonvento & Bolaños, 2021; Wolfe et al., 2024).

To investigate whether *pygl-1* is required in individual neurons with distinct metabolic profiles, we first sought to identify individual neurons with distinct glycolytic states. It had been shown that metabolic states can be predicted based on RNA expression levels of metabolic proteins (Li et al., 2025), so we analyzed single-cell RNA-seq data from the CeNGEN project to identify neurons with differential expression of glycolytic genes (Hammarlund et al., 2018). We observed that ASER sensory neurons showed higher expression of glycolytic genes compared to AIY interneurons (Figure 3-S1), suggesting underlying metabolic heterogeneity between these cell types. To test whether these transcriptomic differences correspond to functional metabolic states, we expressed the HYlight biosensor in ASER (using the *flp-6* promoter) and AIY (using the *ttx-3* promoter) and found that ASER neurons exhibited significantly higher baseline FBP levels than AIY neurons (Figure 3-S2), consistent with the differences in expression of glycolytic genes. We also observed that upon transient hypoxia, both neurons increased their FBP levels, consistent with our previous observations that neurons dynamically regulate their metabolic states upon energy stress (Figure 3-S2) (Wolfe et al., 2024).

**Figure 3:**
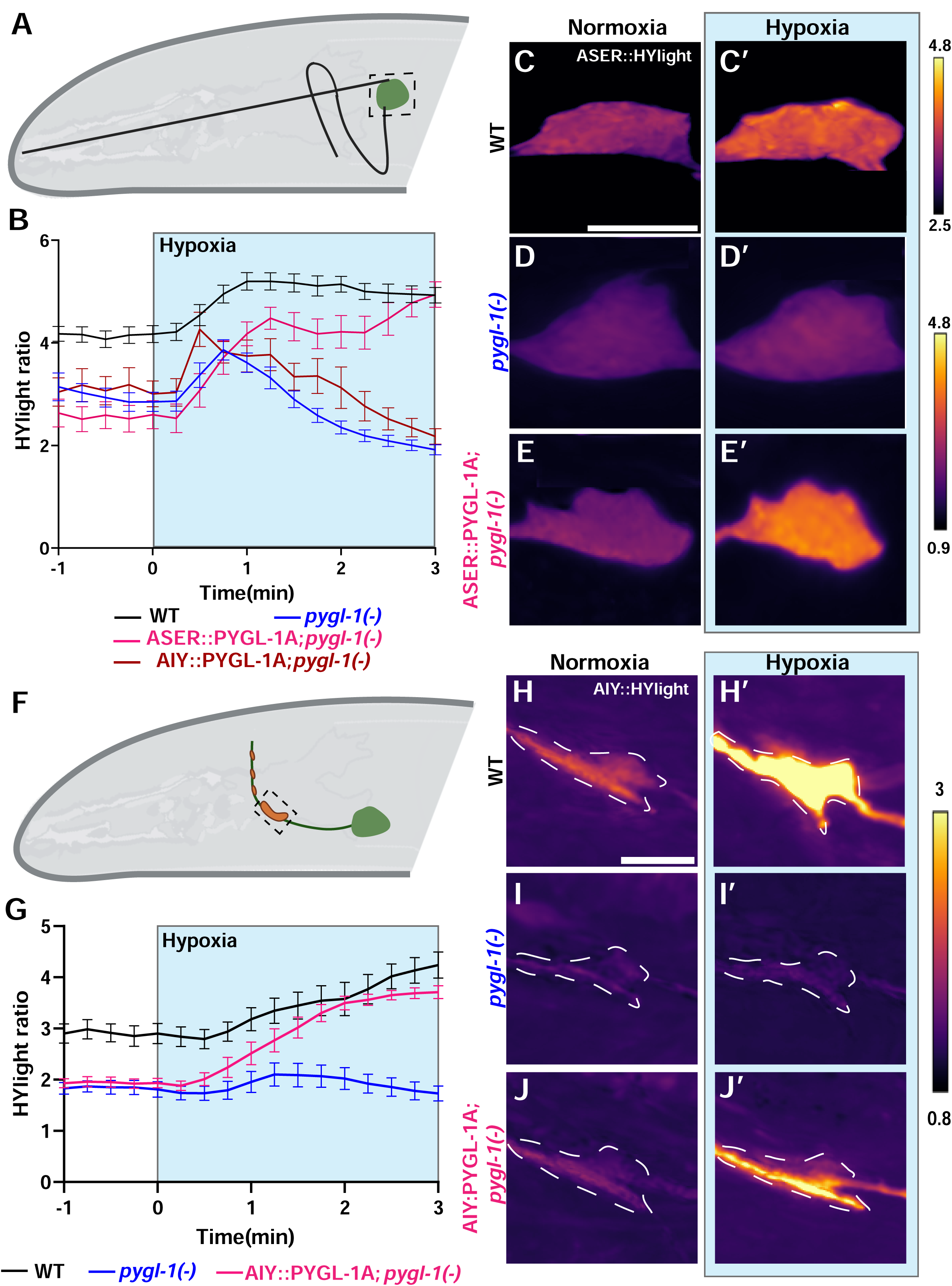
PYGL-1A/Glycogen phosphorylase acts cell-autonomously in neurons. **(A)** Schematic of the ASER sensory neuron located in the head of the worm, with dashed box around the cell body that was imaged in this study. **(B)** Plot of HYlight ratios over time for WT, *pygl-1(tm5211)* mutants, AIY and ASER-specific rescue of PYGL-1A in *pygl-1(tm5211)* mutants. The blue shaded area indicates the period of hypoxia, initiated after 1 minute of normoxia. Error bars represent the standard error of the mean at each time point. Twenty-three animals were analyzed for WT, 19 for *pygl-1(tm5211)*, 8 for AIY specific rescue and 13 for ASER-specific rescue. Note that expressing PYGL-1A in AIY did not rescue FBP levels in ASER, further indicating that PYGL-1A is required cell-autonomously to support glycolytic responses during hypoxic stress. **(C–E)** Ratiometric images of HYlight in ASER of WT worms (C), *pygl-1(tm5211)* mutants (D) and ASER-specific PYGL-1A rescue in *pygl-1(tm5211)* mutants (E). **(C’–E’)** As in (C-E), but after 3 minutes of transient hypoxia. The calibration bars to the right indicates ratiometric signal intensity. In this part of the figure we used a range of 2.5–4.8 for WT and a range of 0.9–4.8 for *pygl-1(tm5211)* mutants and ASER:PYGL-1A rescue to better display the observed differences. **(F)** Schematic of the AIY interneuron with a box outlining the specific synaptic region used for HYlight measurements. **(G)** Plot of HYlight ratios over time for WT, *pygl-1(tm5211)* mutants, and AIY-specific rescue of PYGL-1A in *pygl-1(tm5211)* mutants at AIY Zone 2. The blue shaded area indicates the hypoxia period following 1 minute of normoxia. Error bars represent the standard error of the mean at each time point. Eleven animals were analyzed per genotype. **(H–J)** Ratiometric images of HYlight in AIY Zone 2 of WT animals (H), *pygl-1(tm5211)* mutants (I), and AIY-specific PYGL-1A rescue in *pygl-1(tm5211)* mutants (J). **(H′–J′)** As in (H–J), but after 3 minutes of transient hypoxia. The calibration bar to the right indicates ratiometric signal intensity. Scale bar in (C) and (H) is 5 µm and corresponds to (C-E’ and H-J’).

We next examined if PYGL-1 was required to support the ability of the neurons to dynamically regulate glycolysis upon energy stress, regardless of their starting glycolytic state. We observed that ASER neurons (Figure 3A), which have high glycolytic states in wild type animals, displayed a reduction of the glycolytic state in *pygl-1(tm5211)* mutants in normoxia, and that their FBP levels failed to increase during transient hypoxia (Figure 3B,3C-C’, and D-D’, also consistent with observations in the nerve ring, see Fig 2). To test whether *pygl-1* functions cell-autonomously in ASER, we expressed the PYGL-1A cDNA specifically in ASER neurons in *pygl-1(tm5211)* mutants. We observed that this neuron-specific rescue restored HYlight dynamics during hypoxia to WT levels (Figure 3B, 3E–E′), confirming that PYGL-1 acts cell-autonomously in ASER neurons to support glycolytic plasticity during transient hypoxia. However, expressing the PYGL-1A isoform in AIY neurons using the AIY-specific *ttx-3G* promoter in *pygl-1(tm5211)* mutants did not restore FBP transients in ASER (Figure 3B), further supporting the cell-autonomous role of *pygl-1* in ASER.

We then assessed whether the AIY neurons, which have a low-glycolytic state, also depend on *pygl-1* for their glycolytic responses (Figure 3F). FBP levels in the AIY synaptic regions of *pygl-1(tm5211)* mutants were lower than those in wild-type animals (Figure 3H–I and 3G). During transient hypoxia, HYlight ratios in the AIY synapses of *pygl-1(tm5211)* mutants failed to sustain FBP levels (Figure 3I’ and 3G). The soma of AIY neurons in *pygl-1* mutants also showed reduced FBP levels compared to wild-type (Figure 3-S2). We next asked whether neuron-specific expression of PYGL-1 could restore the hypoxic response in *pygl-1(tm5211)* mutants. Neuron-specific expression of the PYGL-1A cDNA in AIY using the *ttx-3G* promoter in *pygl-1(tm5211)* mutants rescued FBP transients (Figure 3J–J’ and 3G; Figure 3-S2). These data indicate that PYGL-1 is required within AIY neurons to maintain glycolytic plasticity during energy stress.

Together, these results demonstrate that *pygl-1* functions cell-autonomously in both sensory and interneurons—across cell types with distinct metabolic states and gene expression profiles—to support the maintenance and upregulation of glycolytic flux during transient hypoxia. These findings extend our pan-neuronal observations and establish glycogen-dependent glycolytic plasticity (GDGP) as a cell-autonomous mechanism acting in specific neurons to respond to energy stress.

### Activity-induced glycolytic plasticity depends on glycogenolysis and mitochondrial function

Hypoxia and neuronal stimulation are associated with increases in glycolysis in neurons (Díaz-García et al., 2017; Wolfe et al., 2024). Is glycogen breakdown required for increases of glycolysis during neuronal stimulation? To address this question, we turned to the ASER chemosensory neurons, which respond to decreases in NaCl (Suzuki et al., 2008). Building on previous work (Miller et al., 2005; Albrecht & Bargmann, 2011), we adapted microfluidics devices for imaging live animals and added precise control of buffer salt concentrations using a custom-built combinatorial syringe pump system (Figure 4A, Figure 4-S1 and Movie S3). We verified our paradigm by using a calcium indicator, GCaMP, expressed in ASER and confirmed that the neuron responded to decreases in salt concentrations in our assays, consistent with existing literature (Suzuki et al., 2008) (Figure 4-S2).

**Figure 4:**
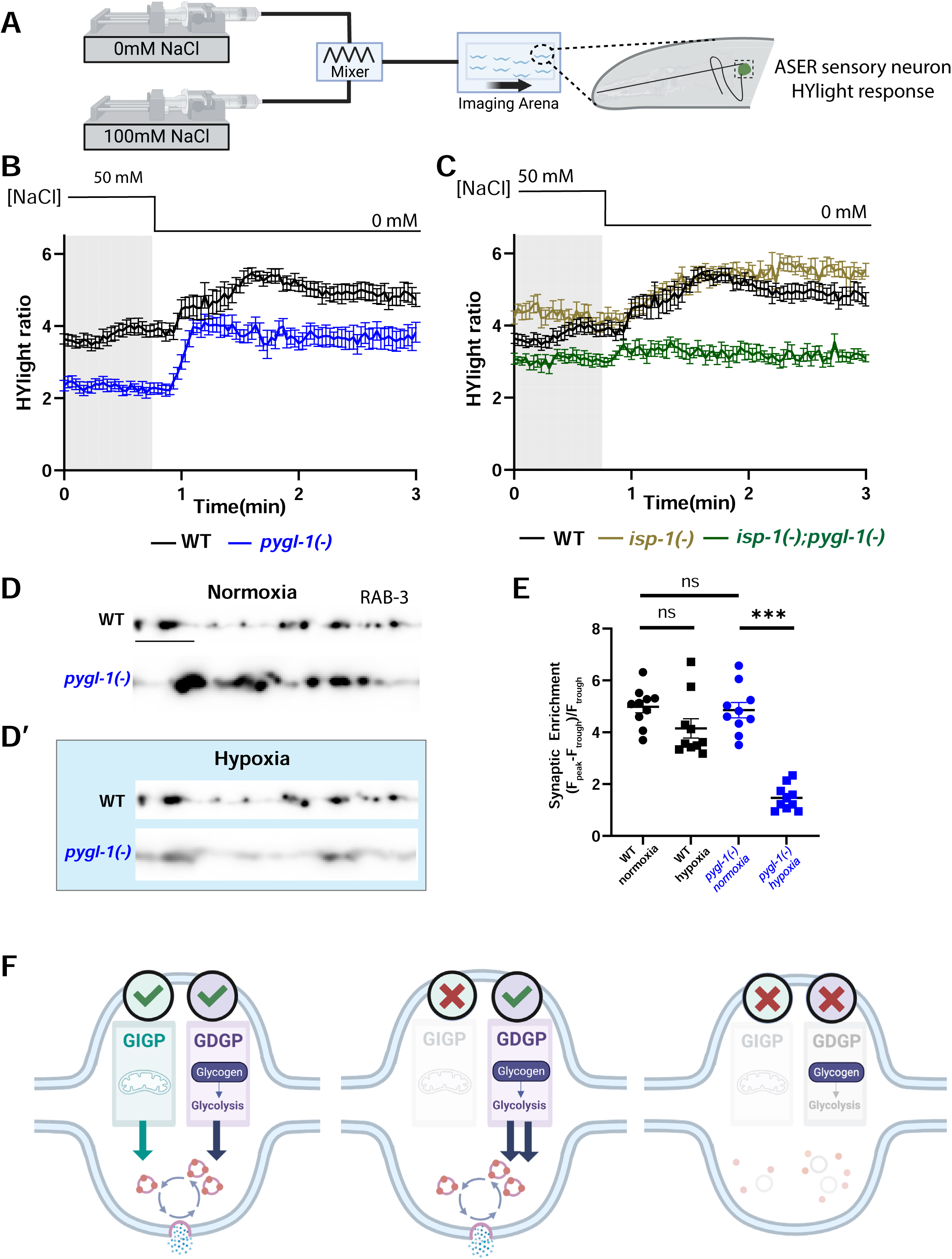
*pygl-1/*Glycogen phosphorylase is required for glycolytic plasticity and synaptic vesicle recycling during mitochondrial impairment. **(A)** Schematic of the dual pump mixing setup and microfluidic device used to deliver desired salt concentrations to a population of worms (see methods) expressing the HYlight biosensor in the ASER neuron. The dashed black box outlines the ASER soma, which was used to measure HYlight ratios. **(B)** HYlight ratios in the ASER soma for WT and *pygl-1(tm5211)* mutants. *pygl-1(tm5211)* mutants have lower baseline values for HYlight than wild type animals (as in Fig 3B and G), but sensory stimulation (using a salt gradient and depicted in line above the graph), *pygl-1(tm5211)* mutants increased FBP levels similarly to WT animals. Animals were stimulated by decreasing the salt concentration from 50 mM to 0 mM NaCl, as shown in the schematic above each graph. The gray shaded area indicates exposure to 50 mM NaCl, and the white region indicates the switch to 0 mM NaCl. Thirteen animals were analyzed for each genotype. **(C)** As in (B), but for WT, *isp-1(qm150)*, and *isp-1(qm150);pygl-1(tm5211)* double mutants. *isp-1(qm150)* mutants, which have defects in mitochondrial function, displayed increases in FBP upon neuronal sensory stimulation similar to wild type. However, the *isp-1(qm150);pygl-1(tm5211)* double mutants did not, phenocopying the *pygl-1(tm5211)* in transient hypoxia and supporting that glycogen breakdown is required for glycolytic plasticity during mitochondrial dysfunction, both in *isp-1* mutants and in transient hypoxia. Thirteen animals were analyzed for WT, nine for *isp-1(qm150)*, and eleven for *isp-1(qm150);pygl-1(tm5211)*. Error bars represent the standard error of the mean ratio at each time point. **(D)** Confocal micrographs representing a Zone 3 neurite during normoxia in WT (above) and *pygl-1(tm5211)* mutants (below). The punctate structures represent vesicle pools in the synaptic region, labeled with synaptic vesicle associated protein RAB-3 tagged with mCherry. (**D’)** Same animal as in (D) but after 10 minutes of transient hypoxia. Note that RAB-3 becomes diffusely localized during transient hypoxia in *pygl-1(tm5211)* mutants, consistent with a defect in synaptic vesicle recycling (Jang et al., 2016). **(E)** Quantification of synaptic enrichment for WT and *pygl-1(tm5211)* mutants (under normoxia and transient hypoxia) as described previously (Jang et al., 2016), see Methods). Each dot corresponds to one worm; ***denotes a p-value of <0.001, as calculated by the Unpaired t-test with Welch’s correction. 10 animals were analyzed per genotype. **(F)** Model summarizing the findings of this study. Neurons employ both glycogen-dependent glycolytic plasticity (GDGP) and glycogen-independent glycolytic plasticity (GIGP) to adapt to changes in energetic demand. GDGP is preferentially utilized under conditions of mitochondrial dysfunction, such as transient hypoxia, and is essential for sustaining glycolytic flux and synaptic vesicle recycling. In contrast, GIGP enables neurons to increase glycolytic output in response to stimuli like neuronal activity, even in the absence of glycogen metabolism, provided mitochondrial function is intact. In the schematic, red dots represent RAB-3 associated with synaptic vesicles. Under conditions of impaired mitochondrial and glycogen metabolism, GDGP cannot be deployed; RAB-3 dissociates from synaptic vesicles and becomes diffusely localized in the cytoplasm, indicating a defect in synaptic vesicle recycling. Scale bar in D is 5µm. Scale bar in D corresponds to (D-D’).

We next examined FBP levels in wild-type ASER cells upon salt-induced neuronal stimulation. Because ASER responds to decreases in salt, we stimulated this neuron with a change from 50 mM to 0 mM NaCl, and observed a concomitant increase in FBP levels (Figure 4B). This FBP response occurred even when the animals were challenged with multiple rounds of salt reduction (Figure 4-S2). Our findings are consistent, and extend, prior literature which demonstrated that increases in neuronal activity induces neuronal glycolysis (Díaz-García et al., 2017).

To determine the role of glycogen metabolism during activity-induced glycolysis, we then assessed FBP levels in the *pygl-1(tm5211)* mutant background. Upon salt-induced neuronal stimulation, *pygl-1(tm5211)* mutants exhibited increases in FBP levels similar to WT animals, despite starting at lower baseline levels (Figure 4B). To investigate whether repeated stimulation would exhaust FBP production in *pygl-1(tm5211)* mutants, we measured HYlight responses following multiple salt pulses. However, under the conditions used in this study, *pygl-1(tm5211)* mutants maintained their ability to increase FBP levels with each salt pulse, similar to WT animals (Figure 4-S2). This contrasts with our hypoxia experiments, where *pygl-1(tm5211)* animals failed to sustain elevated neuronal FBP levels. These results suggest that the same neuron can differentially utilize glycogen depending on the energy stress.

Under hypoxia, where we see that glycolytic responses rely on glycogen breakdown, mitochondrial function is impaired. This led us to hypothesize that the requirement of glycogen phosphorylase in neurons, and glycogen utilization, might be tied to mitochondria function. To test this hypothesis we examined salt-induced FBP responses in ASER neurons of *isp-1(qm150)* mutants, which disrupt mitochondrial function by impairing Complex III of the electron transport chain (J. Feng et al., 2001). We first looked at glycolytic responses in *isp-1(qm150)* mutants, and observed that these mutants maintained glycolytic responses during salt-mediated ASER activity (Figure 4C). Next, we examined the combined effects of blocking glycogen metabolism and mitochondrial respiration by generating *isp-1 (qm150);pygl-1(tm5211)* double mutants. We observed that *isp-1(qm150);pygl-1 (tm5211)* mutants failed to maintain glycolytic plasticity during neuronal activity in ASER neurons (Figure 4C). Together, these findings support a model whereby glycogen-dependent glycolytic plasticity is particularly necessary under conditions of mitochondrial dysfunction, as those seen during transient hypoxia or in *isp-1(qm150)* mutants. Furthermore, it supports a model whereby glycogen-dependent and glycogen-independent glycolytic plasticity mechanisms are differentially deployed in specific contexts and depending on mitochondrial function.

### PYGL-1 is required for synaptic vesicle recycling during energy stress

We next investigated whether the inability of *pygl-1(tm5211)* mutants to increase glycolysis in response to hypoxia has physiological consequences. To address this, we investigated synaptic vesicle recycling in AIY neurons. Previous work from our lab demonstrated defects in neuronal synaptic vesicle recycling in glycolytic mutants under energy stress, as evidenced by diffuse localization of synaptic vesicle-associated proteins, such as RAB-3, upon transient hypoxia (Jang et al., 2016)]. Thus we compared fluorescently-tagged RAB-3 localization in AIY neurons before and after hypoxia treatment. During normoxia, both WT and *pygl-1(tm5211)* mutants maintained clustered localization of the synaptic marker in the neurite region (Figure 4D and 4E), consistent with wild type phenotypes for the synaptic vesicle clusters. However, upon transient hypoxia, *pygl-1(tm5211)* mutants displayed a progressively diffuse localization of the synaptic vesicle marker compared to WT animals, similar to previously reported effects in other glycolytic mutants (Figure 4D’,Figure 4E and Movie S4) (Jang et al., 2016) and consistent with defects in the synaptic vesicle cycle due to energy stress. These results suggest that glycogen breakdown contributes to maintaining normal synaptic vesicle recycling during transient hypoxia. Furthermore, these findings indicate that glycogen breakdown is essential for sustaining glycolytic states in neurons under hypoxic conditions and that this requirement has physiological consequences for synaptic vesicle recycling (Figure 4F).

## Discussion

Neurons dynamically regulate their energy metabolic states *in vivo*. Studies using whole-brain imaging have shown that metabolic states correlate with activity states of brain regions, demonstrating that blood flow and glucose uptake increase in response to brain stimulation (Goense & Logothetis, 2008; Gore, 2003). Similar observations linking activity and changes to metabolism have been seen both at the level of circuits (Mann et al., 2021) and in single neurons (Díaz-García et al., 2017; Wolfe et al., 2024), implicating these metabolic adaptations as direct responses to the changing energetic demands that occur upon neuronal activity. These adaptations are essential for meeting the needs of active synapses, and their impairment can disrupt synaptic functions such as vesicle recycling (Rangaraju et al., 2014; Pathak et al., 2015; Jang et al., 2016; Ashrafi et al., 2020b; Kokotos et al., 2024). While oxidative phosphorylation and glycolysis both respond to changing energetic demands (Ashrafi et al., 2017, 2020a; Díaz-García et al., 2021; Meyer et al., 2022), it remains unclear how and when different fuel sources for these two pathways may be preferentially utilized under specific contexts. In our study we used the HYlight biosensor, which detects the central glycolytic metabolite fructose 1,6-bisphosphate (FBP), to specifically measure glycolysis in single neurons *in vivo* (Koberstein et al., 2022; Wolfe et al., 2024). We observed two cell-specific characteristics of glycolytic metabolism across neuronal classes: 1) neurons *in vivo* can dynamically adjust their glycolytic states in response to energy stress, including during transient hypoxia and changes in neuronal activity; and 2) single neuron classes with distinct identities maintain characteristic baseline HYlight levels, suggesting a diversity of glycolytic metabolic states that map onto neuronal identities in the living organism. Glycogen reserves are a key fuel source for glycolytic metabolism, and we found that neuronal glycogen plays a significant role in supporting adaptive regulation of glycolysis during energy metabolic stress. Our findings indicate that glycogen is important for the synaptic vesicle cycle, suggesting a functional link between glycogen metabolism in neurons and neuronal function. We speculate that this ability of neurons to dynamically adjust energy demands might interface with their capacity to power plastic changes in synaptic function. Together, our results support the emerging idea that energy metabolism is a key axis of neuronal metabolic plasticity and demonstrate that specific fuel sources, like glycogen, contribute to this plasticity under specific contexts.

Glycogen phosphorylase, an enzyme specifically required for glycogen breakdown, is cell-autonomously required in neurons to sustain and support changes in glycolysis The relationship between fuel choice and function is well characterized in non-neuronal tissues such as muscles, where glucose, glycogen, triglycerides, and amino acids fuel metabolic demands and influence physiological functions such as endurance and fatigue resistance (Hargreaves, 2000; Hargreaves & Spriet, 2020; Melzer, 2011). Similarly, brain fuel sources can have distinct physiological effects; for example, a ketogenic diet, which shifts metabolism from glucose to ketones, significantly reduces seizures in drug-resistant epilepsy, likely by decreasing neuronal hyperexcitability (Jensen et al., 2020; Neal et al., 2008). Yet the mechanisms governing neuronal fuel selection during rest and activity remain poorly understood and are subject to ongoing debate. The Astrocyte-Neuron Lactate Shuttle (ANLS) hypothesis suggests that astrocytes metabolize glycogen and export lactate to neurons as an energy source during high activity, implying limited direct glycogen use in neurons (Bélanger et al., 2011; Magistretti & Allaman, 2015). Other studies have demonstrated that glycogen synthase is important to prevent hypoxia-induced cell death in neurons (Saez et al., 2014). Moreover, mice lacking neuronal glycogen exhibit deficits in long-term potentiation (LTP) and learning, suggesting a neuron-specific role for glycogen in cognitive function (Duran, Gruart, Varea, et al., 2019). Our study builds on previous findings by demonstrating that PYGL-1, the enzyme responsible for glycogen breakdown, is required for maintaining glycolytic plasticity in neurons. Specifically, we show that PYGL-1-mediated glycogen metabolism is necessary for sustaining FBP levels during transient hypoxia, supporting the idea that glycogen-derived glucose serves as an important glycolytic fuel in these conditions. Our genetic findings refine and expand current understandings of neuronal glycogen metabolism, and raise important questions regarding the subcellular localization of glycogen and functional compartmentalization in neurons.

Glycogen phosphorylase is specifically required for glycolytic plasticity under conditions of mitochondrial dysfunction. Given the varying energy demands of neurons, metabolic plasticity is essential for adaptation and meeting transient energy demands (Watts et al., 2018). The brain relies on different fuel sources depending on context, and glycogen has previously been shown to support brain metabolism during hypoxia (Brown, 2004; Dienel, 2012; Yellen, 2018). In our study, we observed that glycogen influences glycolytic plasticity through at least two distinct mechanisms acting under specific contexts: glycogen-dependent glycolytic plasticity (GDGP) and glycogen-independent glycolytic plasticity (GIGP). Our findings indicate that GDGP is primarily employed in conditions of mitochondrial dysfunction, such as transient hypoxia or in mutants for mitochondrial function. Why? Under conditions of mitochondrial impairment, energy production will be reduced, and in that context glycogen might act as a capacitor, similar to how it does in muscles (Hargreaves & Spriet, 2020). We speculate that the use of GDGP could be due to the fact that, unlike other energy producing pathways, glycogen does not require ATP input to produce the glycolytic intermediate glucose-6-phosphate (G6P). This is different than the use of glucose, the use of complex sugars (like trehalose), or the use of triglycerides, all which require ATP input to generate intermediates (Judge & Dodd, 2020). The ATP-independent provision of G6P from glycogen may then enable cells to maintain energy production during mitochondrial absence, overload or impairment.

The hypothesis of glycogen as a capacitor in neurons, while speculative, is consistent with our understanding of the role of glycogen in other cells. Studies in yeast highlight glycogen’s role as a metabolic buffer. The glycogen shunt (conversion of G6P to glycogen, as shown in Fig. 1B’) facilitates rapid recovery after glucose replenishment, and disruptions in glycogen synthesis lead to ATP depletion and yeast cell death (Shulman & Rothman, 2017). Our findings indicate that inhibition of glycogen metabolism in neurons results in reduced glycolytic capacity and impaired synaptic function, reinforcing the idea that glycogen serves as a metabolic reserve in neurons and is required as a fuel source under specific contexts. We propose that glycogen functions as an “energy capacitor” that helps buffer changes in energy demands, analogous to how it has been proposed to work in muscles and how a capacitor stabilizes electrical supply during fluctuations in input(Hargreaves & Spriet, 2020). Our findings also highlight the metabolic flexibility of neurons via the context-specific utilization of glycogen.

## Methods

### *C. elegans* culturing

Nematodes were cultured on nematode growth media (NGM) plates at 20°C or 15°C, with OP50 Escherichia coli as their food source. For the wild-type strain, C. elegans Bristol N2 was used. Plates were seeded with OP50 bacteria, and 3-4 worms were transferred onto fresh plates every 3-4 days. Worms fed for at least 2-3 generations were used for experiments. One day adult worms were used for all experiments. The strains utilized in this report can be found in Supplementary Table S1.

### Molecular Biology and Transgenics

We utilized Gibson Assembly or the Gateway system to construct plasmids for generating transgenic *C. elegans* strains. Transgenic strains were produced through standard microinjection techniques, with the construct of interest injected at concentrations ranging from 5 to 100 ng/μL. The following co-injection markers were used-*unc-122*p::GFP, *unc-122*p::RFP, *elt7*::mCHNLS,*elt7*::GFPNLS. The *ttx-3* [specifically *ttx-3* based promoter G], *flp-6* promoters were used to express genetic sequences in AIY and ASE. *col-19* promoter was used to express proteins in the hypodermis. The PYGL-1A cDNA construct was synthesized by amplifying from a cDNA pool derived from a mixed population of *C. elegans*.

CRISPR-Cas9 was used to generate the *pygl-1(ola587)* genetic knockout allele based on earlier protocols (Dickinson & Goldstein, 2016). The CRISPOR tool (Concordet & Haeussler, 2018) was used to select gRNA sites adjacent to the 5′ and 3′ ends of the *pygl-1* coding region, and the corresponding crRNAs were ordered from Horizon Discovery. CRISPR editing was performed by injecting 3 μM Cas9 protein (Horizon) preincubated with tracrRNA (40 μM, Horizon) at 37 °C for 10 minutes. Both crRNAs (30 μM each) and a ∼150-base single-stranded DNA repair template (0.5 μM, Keck Oligo Core, gel-purified) were included in the injection mix. The repair template was designed to delete the *pygl-1* locus via 30-base homology arms flanking the target region. The injection buffer also included 25 mM KCl and 7.5 mM HEPES, pH 7.3. The mix contained additional crRNA (12 μM) and a template (0.25 μM) for recreating the *dpy-10(cn64)* allele as a co-CRISPR marker. F1 progeny exhibiting the roller phenotype were singled and genotyped by PCR and sequencing to confirm the successful generation of the *pygl-1(ola587)* deletion allele.

### Microfluidics and Mounting

For hypoxia imaging, the worms were imaged using a PDMS microfluidic device that enabled control of the atmospheric conditions, as previously described (Jang et al., 2021; Wolfe et al., 2024). In short, a 10% agarose dissolved in water was applied to the top of the PDMS device. Depending on the imaging, a 3μL drop of M9 buffer, or 10 mM levamisole in M9, was placed onto the pad. Subsequently, worms were transferred into the center of the drop, and a size 1.5 coverslip was gently lowered onto the pad. One can also use 2-3% agar for such experiments, but worms might move more.

For salt-mediated ASER activity imaging, a different microfluidic system was used, building upon previous work (Albrecht & Bargmann, 2011; Miller et al., 2005). Two NE-1000 syringe pumps (New Era Pump Systems, Inc; Farmingdale, NY, USA) were loaded with buffers simulating NGM conditions (25 mM K(H)PO4 pH 6, 1 mM MgSO4, 1 mM CaCl2, 5 mM levamisole); one pump was designated NGM.0 (with 0 mM NaCl) and the other NGM.100 (100 mM NaCl). Glycerol was added to adjust to a final osmolality of 260 mOsm for both buffers. These were proportionally combined with a 600 µm diameter passive herringbone microfluidics mixer (Darwin Microfluidics, Inc; Paris, France) to combine the two streams into a single flow containing a desired concentration of NaCl. Custom control software was written in python for delivering desired pumping protocols to the pumps. For experiments, we adapted the P10 device previously described (Albrecht & Bargmann, 2011), but by punching only a single inlet, outlet, and worm chamber entry port; the assembled device was vacuum degassed by clamping together with a 24×50 mm size 1.5 coverslip using a PH-2 clamp from Warner Instruments (Holliston, MA, USA) The device inlet was then connected to the outlet of the mixer, while the exit port was connected to a waste container. The device was then filled with NGM.50 solution using the pumps, and the worms were transferred into the main arena using a syringe. Worms were maintained in the device for 15 minutes prior to an experiment to allow for temperature equilibration and for the levamisole to paralyze the worms.

### Imaging

Microscopy was conducted using a Nikon Ti2 + CSU-W1 spinning disk confocal microscope. Images were captured using a Hamamatsu Orca-Fusion BT CMOS camera at 16-bit pixel depth. Samples were excited with 50 mW lasers at 405, 488, or 561 nm wavelengths, depending on the requirements. For ratiometric imaging of HYlight and GCaMP, the 488 and 405 nm lasers were alternated without changing the 525 nm emission filter. The laser power (488,405 nm respectively) was set to either 8%/2% (hypodermal imaging) or 48%/4% (single neuron/panneuronal imaging) for 10× or 4X objective imaging and 24%/4% for 60× imaging[panneuronal], with the same exposure time (10 to 300 ms). For GCaMP experiments, the laser power (488,405 nm respectively) was set to 32%/8% for both channels with an exposure time of (150-300)ms. The dsRED channel was used to identify ASER while imaging the ASE neurons using a laser power of 75% with 50ms exposure. The exposure time was adjusted for different sample brightness but kept equal across both channels. During hypoxia experiments, the flow was manually switched between tanks of compressed air or pure nitrogen gas, as indicated by the times on the charts. Similarly, for the ASER activity experiments, the salt buffer flow was combined (using a mixer) or switched between NGM buffer containing 100 mM NaCl or 0 mM NaCl using single channel pumps as indicated on the charts. The flow rate was set to 0.4 ml/min for all experiments. For synaptic enrichment quantification, images were captured using a 60× objective with the 561 nm laser at 40% power and 200 ms exposure time.

### Feeding RNAi

The MRC RNAi library was used to perform feeding RNAi experiments (Horn et al., 2010; Kamath et al., 2003). The RNAi clones were grown overnight in LB media containing ampicillin/carbenicillin (diluted 1:1000 from 100mg/ml stock). This bacteria was seeded onto NGM plates containing IPTG to a final concentration of 1mM and carbenicillin to a final conc of 25μg/ml. Note: IPTG and carbenicillin were added after the autoclave step of making the NGM plates. All other steps were followed as per standard NGM plate making protocols. The plates were left to dry for a day at room temperature and then worms were transferred onto the plates as L4s/adults. Worms were fed for 1 generation before imaging.

### Data Analysis

#### Comparison of glycolytic gene RNA expression data

Binary expression patterns of single-cell RNA-seq data for all neurons of *C. elegans* animals at the L4 stage (Taylor et al., 2021) was downloaded from www.cengen.org at the Threshold 2 level. The log of the normalized TPM values +1 from this file were then calculated for each gene in the glycolytic pathway as shown in Figure 3 S1 for all neurons. The mean and standard deviation for these values were then used to calculate a Z score for each neuron based on the relative enrichment relative to all neurons; these Z scores were then compared for AIY and ASER. Genes that were filtered at Threshold 2 for both neurons were excluded from analysis.

#### HYlight quantification

Images were processed using Fiji/ImageJ software. For visualization, ratiometric images were generated by dividing the 488-nm excitation channel by the 405-nm channel. The LUT was set to mpl-inferno and scaled to the ratio ranges per figure. Note-Neurons and the hypodermis were imaged at different laser settings. Therefore we cannot compare raw HYlight ratios between those different tissues.

<u>Neuronal imaging:</u> Quantification of 10X and 60X HYlight images was similar to previously described protocol (Wolfe et al., 2024). Briefly, for 60X (Panneuronal imaging) we used scripts that applied background subtraction with a Gaussian filter to both channels and then thresholded to generate an ROI based on the threshold. The resulting ROI was then used to capture pixel values from the corresponding ratiometric image slices. These pixel values were exported and analyzed using GraphPad Prism. For 4x (ASER salt activity imaging) or 10× (AIY or ASER hypoxia imaging) captures, background subtraction was applied, and an ROI was drawn around the region of interest (typically the cell soma or neurite) per time frame. The mean pixel value of this ROI was used as the representative value for each cell.

<u>Hypodermis imaging:</u> Hypodermal HYlight was captured using a 10× objective and processed using Fiji/ImageJ similar to above. The ROI was drawn in the hyp 7 cell of the hypodermis, below/around the worm pharynx region, and the mean pixel value was used as the representative value for each worm. Overall, in HYlight graphs, One way ANOVA was used for statistical analyses with more than 2 comparisons, whereas t-tests were used for 2 data sets [will be mentioned next to the graph]

#### Synaptic Enrichment quantification

To quantify synaptic enrichment of synaptic vesicle proteins, fluorescence values for individual neurites, Zone 3 for the AIY neuron were obtained through segmented line scans using ImageJ. A sliding window of 2μm was used to identify all the local fluorescence peak values and trough values for an individual neuron (the maximum and the minimum fluorescence values in a 2μm interval, respectively). Synaptic enrichment was then calculated as %ΔF/F as previously described (Jang et al., 2016). Briefly, all the identified local maximum and minimum fluorescence values in a given neurite (local Fpeak and local Ftrough) were averaged and used to calculate ΔF/F, with ΔF/F being the difference between average peak-to-trough fluorescence (F) defined as (Fpeak − Ftrough)/Ftrough. One-way ANOVA was used for statistical analyses.

## Acknowledgements

We thank current and former members of the D.C.-R. lab for advice, suggestions, and guidance, in particular Snusha Ravikumar. We thank the Caenorhabditis Genetics Center (P40 OD010440) and the Mitani lab (Tokyo Women’s Medical University School of Medicine) for strains. We thank the Keck Oligo Synthesis Resource at Yale for their assistance with synthesis of short single-stranded DNA oligos and Azenta Life Sciences for help with DNA sequencing. Some figures were created in https://BioRender.com. This work was supported by the grant R35 NS132156 awarded to D.A.C.-R.

