## Supplementary Info for "Glycogen metabolism acts in neurons to support glycolytic plasticity"

**Figure 1-S1: Measurements of HYlight signal in the hypodermis. (A)** A ratiometric image of the HYlight biosensor expressed in hypodermis using the *col-19* promoter shows the emission ratio after 488nm and 405 nm excitation. The white dashed boxes show the regions selected for quantification in (B). (**B)** A plot of HYlight ratios over time for different regions in the hypodermis. The shaded area indicates the period of transient hypoxia after 1 minute of normoxia. 8 animals were used in the analysis per region for all regions except for the tail, in which 7 animals were used. The thick line in the graph represents the region “below the pharynx”, which was selected for further analysis, owing to its high signal-to-noise ratio combined with a robust response to hypoxia. **(C)** A plot of hypodermal HYlight ratios over time for worms fed bacteria with empty vector and *pfk-1.1* RNAi. The shaded area indicates the period of hypoxia after 1 minute of normoxia. Error bars represent the standard error of the mean at each time point, with 10 animals analyzed per strain. Scale bar in (A) is 50µm.

**Figure 1-S2 : RNAi screen used to identify *pygl-1* as a key regulator of glycolysis under normoxia and hypoxia. (A)** Schematic representation of cellular metabolic pathways adjacent to glycolysis examined in this study. Underlined genes were targeted for knockdown by feeding dsRNA to worms. Targeted genes included glycolytic enzymes (*hxk-2*, *pgk-1*), trehalose synthesis (*gob-1*), glycogen breakdown (*pygl-1*), triglyceride breakdown (*atgl-1*, *hosl-1*), mitochondrial respiration (*cco-1*), and transporters (*fgt-1*, *mct-3*). The goal was to identify gene knockdowns that disrupt glycolytic function similarly to *pfk-1.1* RNAi. **(B)** Quantification of hypodermal HYlight ratios during normoxia across RNAi-treated worms. Each dot represents one worm; **** indicates $p < 0.0001$, *** indicates $p < 0.001$, calculated using Brown-Forsythe and Welch’s ANOVA followed by Dunnett’s multiple comparisons test. Ten animals were analyzed per condition. Among the genes tested, only RNAi against *pfk-1.1*, *pygl-1*, and *cco-1* significantly reduced FBP levels compared to empty vector controls. **(C)** Quantification of hypodermal HYlight ratios during hypoxia for worms treated with RNAi targeting *pfk-1.1*, *pygl-1*, and *cco-1*, compared to empty vector controls. Each dot represents one worm; **** indicates $p < 0.0001$, calculated using Brown-Forsythe and Welch’s ANOVA followed by Dunnett’s multiple comparisons test. Ten animals were analyzed per condition. Only *pygl-1* RNAi reduced FBP levels under hypoxia to levels comparable to *pfk-1.1* RNAi, highlighting a role for glycogen breakdown in maintaining glycolytic flux during hypoxic stress.

**Figure 1-S3: Loss of *pygl-1* reduces hypodermal FBP levels during normoxia and hypoxia. (A)** Plot of hypodermal HYlight ratios over time in worms treated with empty vector or *pygl-1* RNAi. The shaded area indicates the hypoxia period, initiated after 1 minute of normoxia. Error bars represent the standard error of the mean at each time point. Ten animals were analyzed per genotype. **(B)** Plot of hypodermal HYlight ratios over time in WT and *pfk-1.1(ola458)* mutant worms. The shaded area indicates the hypoxia period following 1 minute of normoxia. Error bars represent the standard error of the mean at each time point. Ten animals were analyzed per genotype. **(C)** Plot of hypodermal HYlight ratios over time in WT and *pygl-1(tm5211)* mutant worms. The shaded area indicates the hypoxia period following 1 minute of normoxia. Error bars represent the standard error of the mean at each time point. Ten animals were analyzed per genotype. Similar to the RNAi experiments, the *pygl-1(tm5211)* mutation reduced hypodermal FBP levels to levels comparable to those observed in *pfk-1.1* null animals during both normoxia and hypoxia.

**Figure 3-S1: Glycolytic gene expression in ASER sensory neurons and AIY interneurons. (A)** Schematic of the glycolytic pathway indicating genes analyzed; genes shown in gray were excluded due to lack of detectable expression in both neurons. **(B)** Comparison of RNA expression levels for glycolytic genes in ASER sensory neurons (pink) and AIY interneurons (grey), based on single-cell RNA-seq data from the CeNGEN project (Hammarlund et al., 2018). Z-scores were calculated as the log-transformed transcripts per million (TPM) across all neurons, per gene, and represent relative transcript enrichment compared to the neuronal mean.

**Figure 3-S2: AIY interneurons have lower baseline FBP levels than ASER sensory neurons and require PYGL-1 for hypoxia-induced glycolytic responses. (A)** Schematic of the AIY neuron (left) and ASER neuron (right) in the head of the worm. The boxed regions indicate the soma of each neuron, where HYlight ratios were measured for this figure (as compared to the synaptic region in AIY, in Figure 3). **(B)** Comparison of HYlight ratios in the soma of AIY and ASER neurons under normoxia in WT animals. Baseline FBP levels in AIY are significantly lower than in ASER, consistent with the lower expression of glycolytic genes (Figure 3 S1). Each dot represents one worm; *** denotes $p < 0.001$, calculated using an unpaired *t*-test. Error bars represent the standard error of the mean. Ten animals were analyzed per genotype. **(C)** Plot of HYlight ratios over time in AIY and ASER neurons in WT animals. AIY exhibits lower baseline FBP levels than ASER, but both neurons show an increase in FBP during transient hypoxia. Ten animals were analyzed per genotype. **(D)** Plot of AIY soma HYlight ratios over time in WT, *pygl-1(tm5211)* mutants, and AIY-specific rescue of PYGL-1A in *pygl-1(tm5211)* mutants. Rescue in AIY (as shown in A, left schematic) restores the FBP increase during hypoxia in *pygl-1(tm5211)* mutants, consistent with what was observed for the synaptic regions in Figure 3G. Ten animals were analyzed per genotype. In both (C) and (D), the gray shaded area indicates the hypoxia period, which followed 1 minute of normoxia. Error bars represent the standard error of the mean at each time point.

**Figure 4-S1: Validation of the dual-pump microfluidic device used for this study for rapid changes to delivered buffer conditions**. **(A)** A dual-pump system was used for delivering salt concentrations to worms via microfluidics, and the delay and effectiveness of this method was validated by combining a non-fluorescent buffer (Pump A) with a fluorescent buffer (Pump B) together at various proportional combinations, which resulted in linear changes to fluorescence as expected. **(B)** A comparison of the expected protocol for changes in concentration in terms of the percent flow rate of the pump containing the fluorescent dye (Pump B) with the measured fluorescence. Shaded region reflects the standard deviation of five positions within the chamber. Fluorescence buffer is 0.5% Ponceau S, imaged using standard GFP/FITC settings.

**Figure 4-S2: Neuronal-activity induced calcium and glycolytic responses in ASER neurons. (A)** Schematic of the ASER neuron in the head of the worm. The dashed black box surrounds the soma of ASER, where GCaMP and HYLight ratios were measured. **(B)** GCaMP ratios over time in the ASER soma of WT worms. Neuronal stimulation (with salt, stimuli depicted above the graph) dynamically increases calcium levels. Six WT animals were analyzed. **(C)** HYlight ratios over time in the ASER soma of WT and *pygl-1(tm5211)* mutants during multiple pulses of salt stimulation. Eleven animals were analyzed for WT and ten for *pygl-1(tm5211)*. *pygl-1(tm5211)* mutants increased FBP levels similarly to WT animals during repeated salt stimulation. In both (B) and (C), animals were stimulated by decreasing the salt concentration from 50 mM to 0 mM NaCl, as indicated in the schematic above each graph. The gray shaded area represents exposure to 50 mM NaCl, and the white region indicates the switch to 0 mM NaCl. Error bars represent the standard error of the mean at each time point.

**Table S1:** List of strains used in this study. Note: CGC is the Caenorhabditis Genetics Center, NBRP is the National Bioresource Project (Strains from Shohei Mitani).

| Strain | Genotype | Source |
|---|---|---|
| N2 | Bristol wild-type strain | CGC |
| DCR9288 and DCR9299 | *olaIs141* and *olaIs142* [Both are *flp-6*::*HYlight* strains] | This study |
| DCR 9685 | *isp-1(qm150); pygl-1(tm5211)/tmc16; olaIs142* | This study |
| DCR9573 | *olaIs151* [*rab-3p*::*HYlight*] | This study. The non-integrated extrachromosomal array strain from (Wolfe et al., 2024) |
| DCR9683 | *pfk-1.1(ola458); olaIs151* | This study. The non-integrated extrachromosomal array strain from (Wolfe et al., 2024) |
| DCR9890 | *pygl-1(ola587);olaIs151* | This study |
| DCR9861 | *pygl-1(tm5211); olaEx5694; olaIs151* [*olaEx5694* is *rab-3p::pygl-1a*] | This study |
| DCR9717 | *olaIs123 [ttx-3p::rab-3::mCh*] | This study |
| DCR9666 | *pygl-1(tm5211); olaIs123* | This study |
| DCR9656 | *isp-1(qm150); olais141* | This study |
| DCR9640 | *pygl-1(tm5211)/tmc16; olaIs151* | This study |
| DCR9639 | *pygl-1(tm5211)/tmc16; olais142* | This study |
| DCR9229 | *olaEx5472 [olaEx5472 is flp-6::GCaMP8m*] | This study |
| DCR9886 | *pygl-1(tm5211); olaEx5702; olaIs142* [*olaEx5702* is *flp-6p::pygl-1a*] | This study |
| *DCR9089* | *olaIs138 [ttx-3p::HYlight]* | (Wolfe et al., 2024) |
| DCR9565 | *pygl-1(tm5211); olaIs138* | This study |
| DCR9568 | *pygl-1(tm5211); olaEx5653; olaIs138* [*olaex5653* is *ttx-3p::pygl-1a*] | This study |
| DCR9191 | *olaEx5464 [olaEx5464* is *col-19p::HYlight]* | This study |
| DCR9459 | *pygl-1(tm5211)/tmc16; olaEx5464* | This study |
| DCR9193 | *pfk-1.1(ola458); olaEx5464* | This study |
| tm5211 | *pygl-1(tm5211)* | NBRP |
| FX30161 | *tmC16 [unc-60(tmIs1237)]* | NBRP |

**Movie S1 (separate file): HYlight responses in neurons upon hypoxia in WT animals.** HYlight dynamics during transient hypoxia in neurons of WT worms mounted in M9 buffer on a microfluidic device (described in Methods). Images were captured using two channels in a single *z*-plane every 5 seconds at 4× magnification. Hypoxia was initiated at 0 minutes of imaging, following an initial 1-minute normoxic phase, as indicated by a dot appearing in the top right corner of the video. Upon hypoxia, a global increase in FBP levels is observed across neurons in all worms, consistent with enhanced glycolytic activity. The calibration bar in the top right displays HYlight ratios, with colors representing the magnitude of the ratiometric signal. Scale bar: 100 μm.

**Movie S2 (separate file): Loss of *pygl-1* abolishes neuronal FBP increase during hypoxia.** HYlight dynamics in neurons of *pygl-1(tm5211)* mutant worms mounted in M9 buffer on a microfluidic device (described in Methods). Images were captured using two channels in a single *z*-plane every 5 seconds at 4× magnification. Hypoxia was initiated at 0 minutes of imaging, following an initial 1-minute normoxic phase, as indicated by a dot appearing in the top right corner of the video. Unlike WT animals, *pygl-1(tm5211)* mutants do not exhibit an increase in HYlight ratios during hypoxia, suggesting impaired glycolytic upregulation. The calibration bar in the top right displays HYlight ratios, with colors representing the magnitude of the ratiometric signal. Scale bar: 100 μm.

**Movie S3 (separate file): Rapid concentration changes via proportional pump flow mixing.** Fluorescence image of the microfluidics device used for salt stimulation showing a protocol where the proportion of the second pump (Pump B) begins on 50%, is switched between 25% and 75% three times, and ends back on 50%. Buffer transitions were visualized by loading a solution of 0.5% Ponceau S into the second pump channel and captured using standard GFP excitation and emission settings. See Figure 4 S1.

**Movie S4 (separate file): Loss of *pygl-1* disrupts RAB-3 localization in AIY Zone 3 during hypoxia**. Live imaging of the synaptic vesicle-associated protein RAB-3 in the Zone 3 region of the AIY neuron. The video shows a Zone 3 neurite in a *pygl-1(tm5211)* mutant, where RAB-3 is tagged with mCherry. Punctate structures represent synaptic vesicle-associated RAB-3. Images were captured every 30 seconds at 60× magnification using 0.6 μm z-step intervals. The video shown is a maximum intensity projection of the z-slices. Hypoxia was initiated at 0 minutes of imaging following a 1-minute normoxic phase, as indicated by the timestamp in the top left corner. In *pygl-1(tm5211)* mutants, RAB-3 became diffusely localized during hypoxia, consistent with impaired synaptic vesicle recycling. Scale bar: 5 μm.

Figure 1-S1

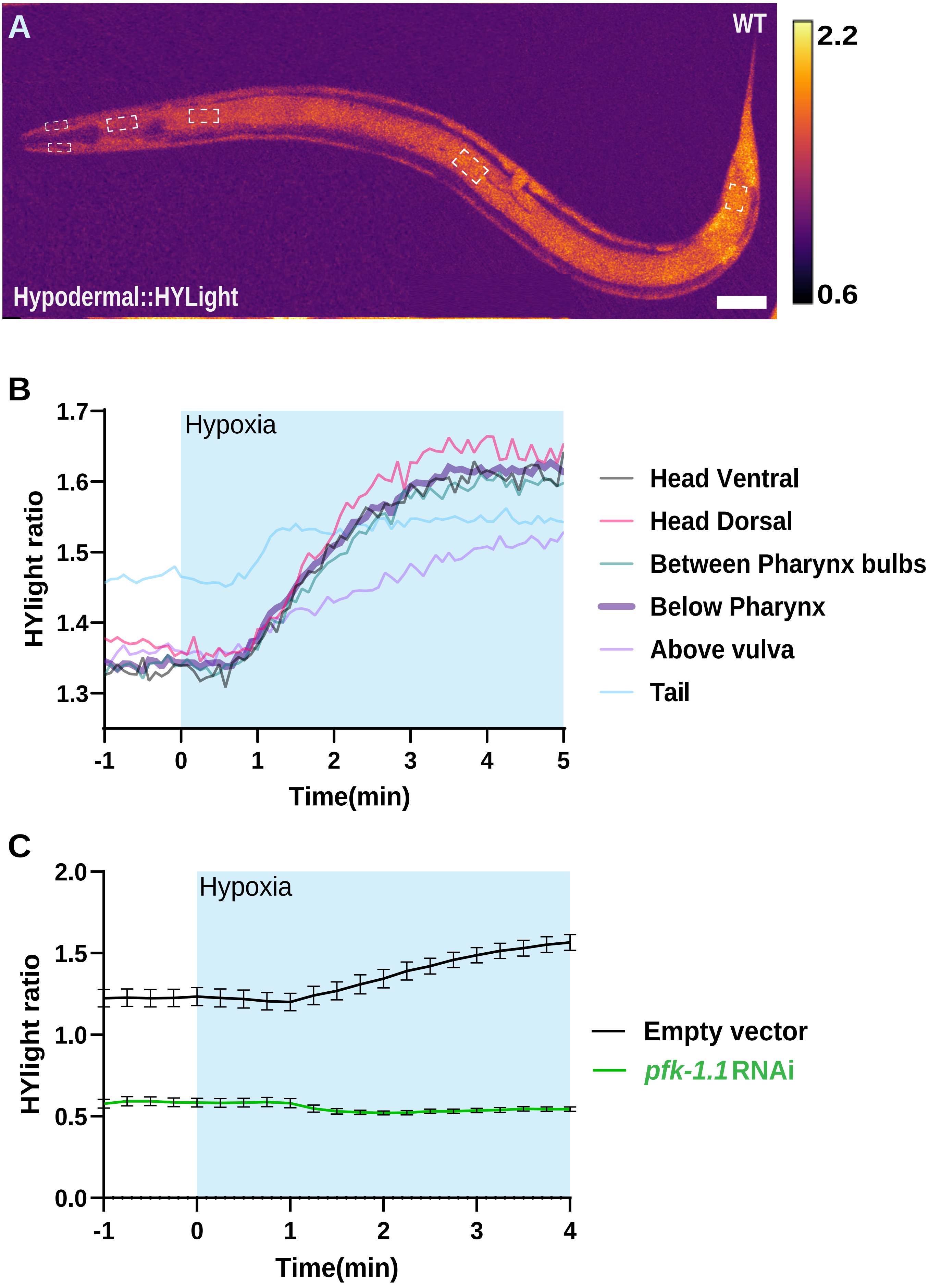
A
WT
2.2
0.6
Hypodermal::HYLight
B
HYlight ratio
1.7
1.6
1.5
1.4
1.3
Hypoxia
-1
0
1
2
3
4
5
Time(min)
Head Ventral
Head Dorsal
Between Pharynx bulbs
Below Pharynx
Above vulva
Tail
C
HYlight ratio
2.0
1.5
1.0
0.5
0.0
Hypoxia
-1
0
1
2
3
4
Time(min)
Empty vector
pfk-1.1 RNAi

Figure 1-S2

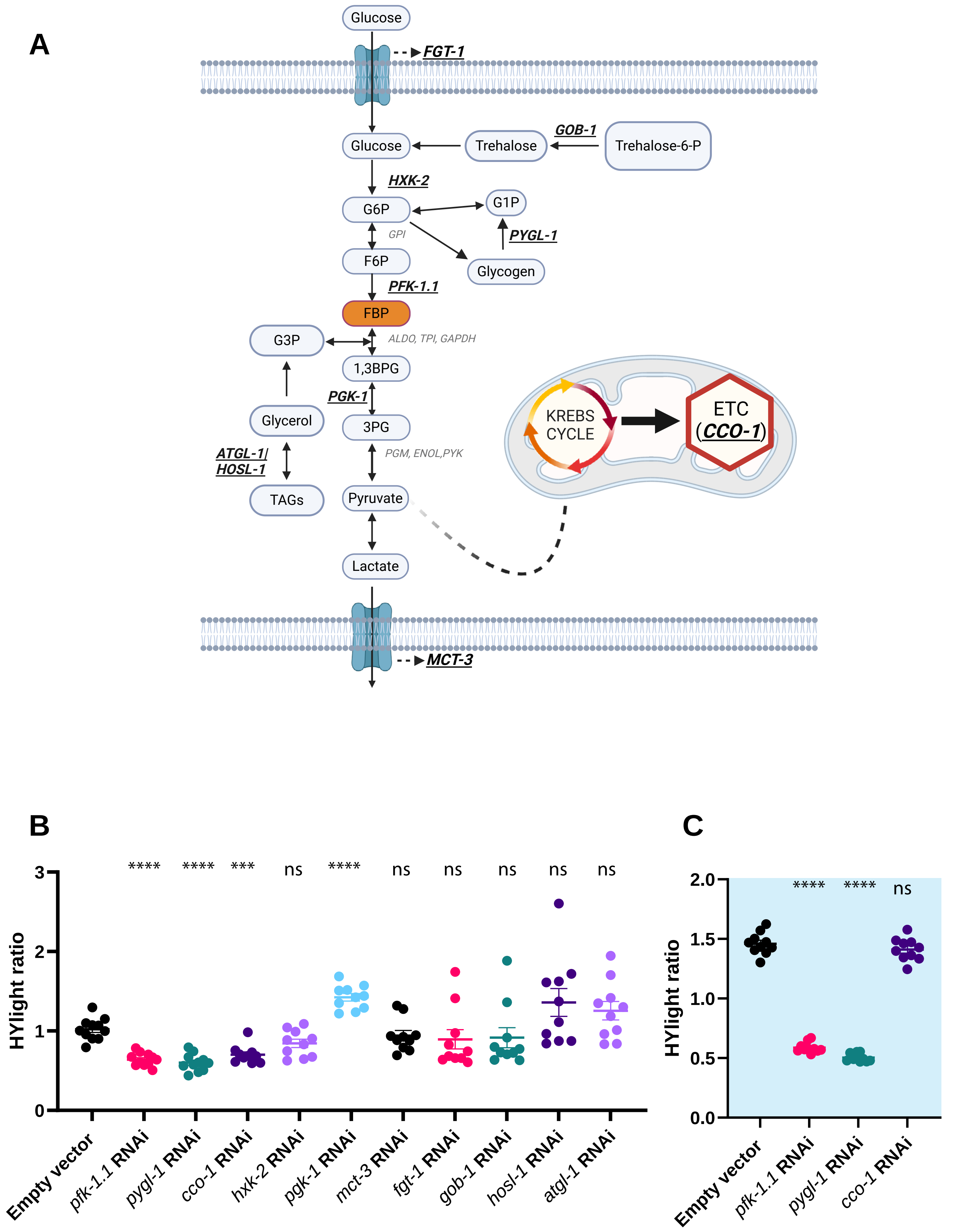

A
Glucose
FGT-1
Glucose
Trehalose
GOB-1
Trehalose-6-P
HXK-2
G6P
G1P
GPI
PYGL-1
F6P
Glycogen
PFK-1.1
FBP
G3P
ALDO, TPI, GAPDH
1,3BPG
PGK-1
Glycerol
3PG
KREBS CYCLE
ETC (CCO-1)
ATGL-1/ HOSL-1
PGM, ENOL,PYK
TAGs
Pyruvate
Lactate
MCT-3
B
HYlight ratio
3
2
1
0
**** **** *** ns **** ns ns ns ns ns
Empty vector
pfk-1.1 RNAi
pygl-1 RNAi
cco-1 RNAi
hxk-2 RNAi
pgk-1 RNAi
mct-3 RNAi
fgt-1 RNAi
gob-1 RNAi
hosl-1 RNAi
atgl-1 RNAi
C
HYlight ratio
2.0
1.5
1.0
0.5
0.0
**** **** ns
Empty vector
pfk-1.1 RNAi
pygl-1 RNAi
cco-1 RNAi

Figure 1-S3

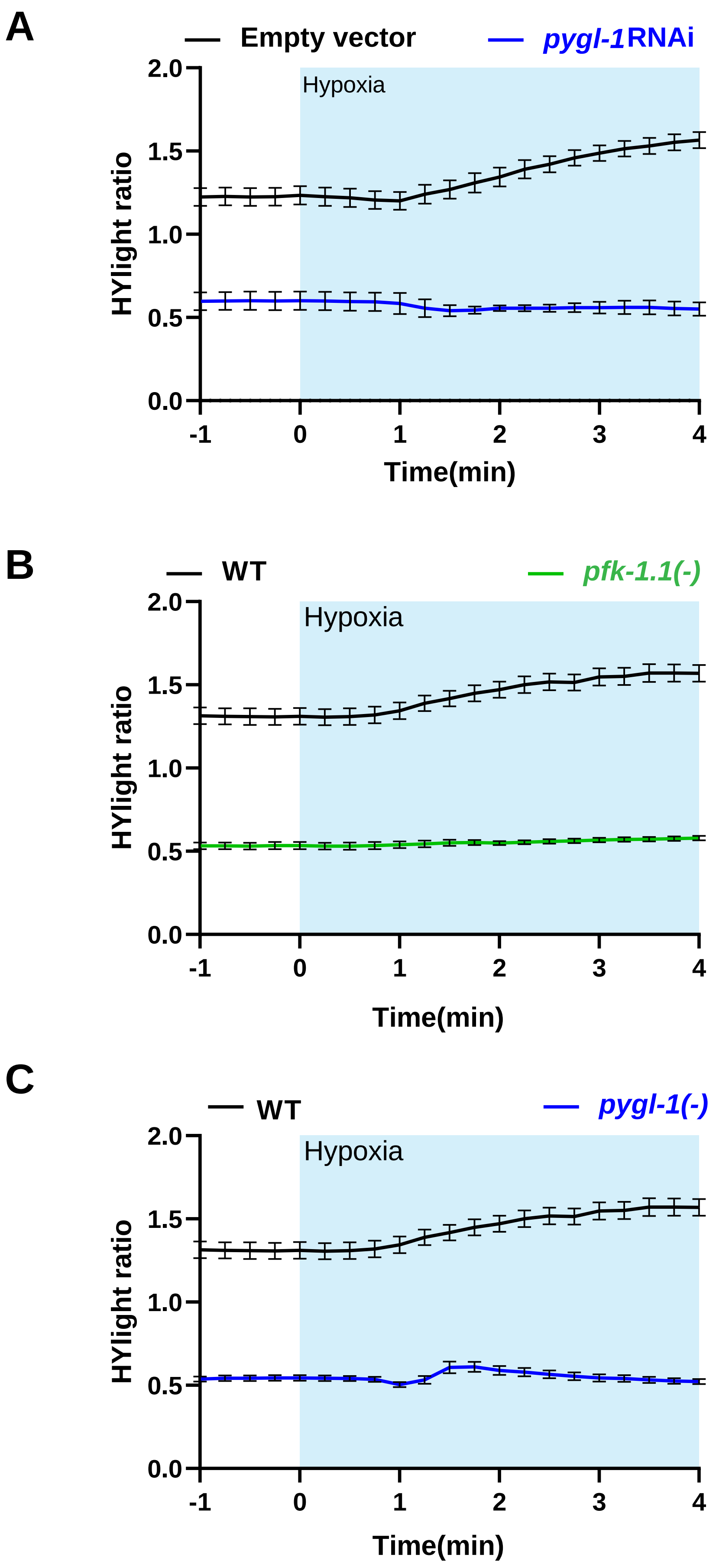

A
Empty vector
pygl-1 RNAi
Hypoxia
HYlight ratio
2.0
1.5
1.0
0.5
0.0
-1
0
1
2
3
4
Time(min)
B
WT
pfk-1.1(-)
Hypoxia
HYlight ratio
Time(min)
C
WT
pygl-1(-)
Hypoxia
HYlight ratio
Time(min)

### Figure 3-S1

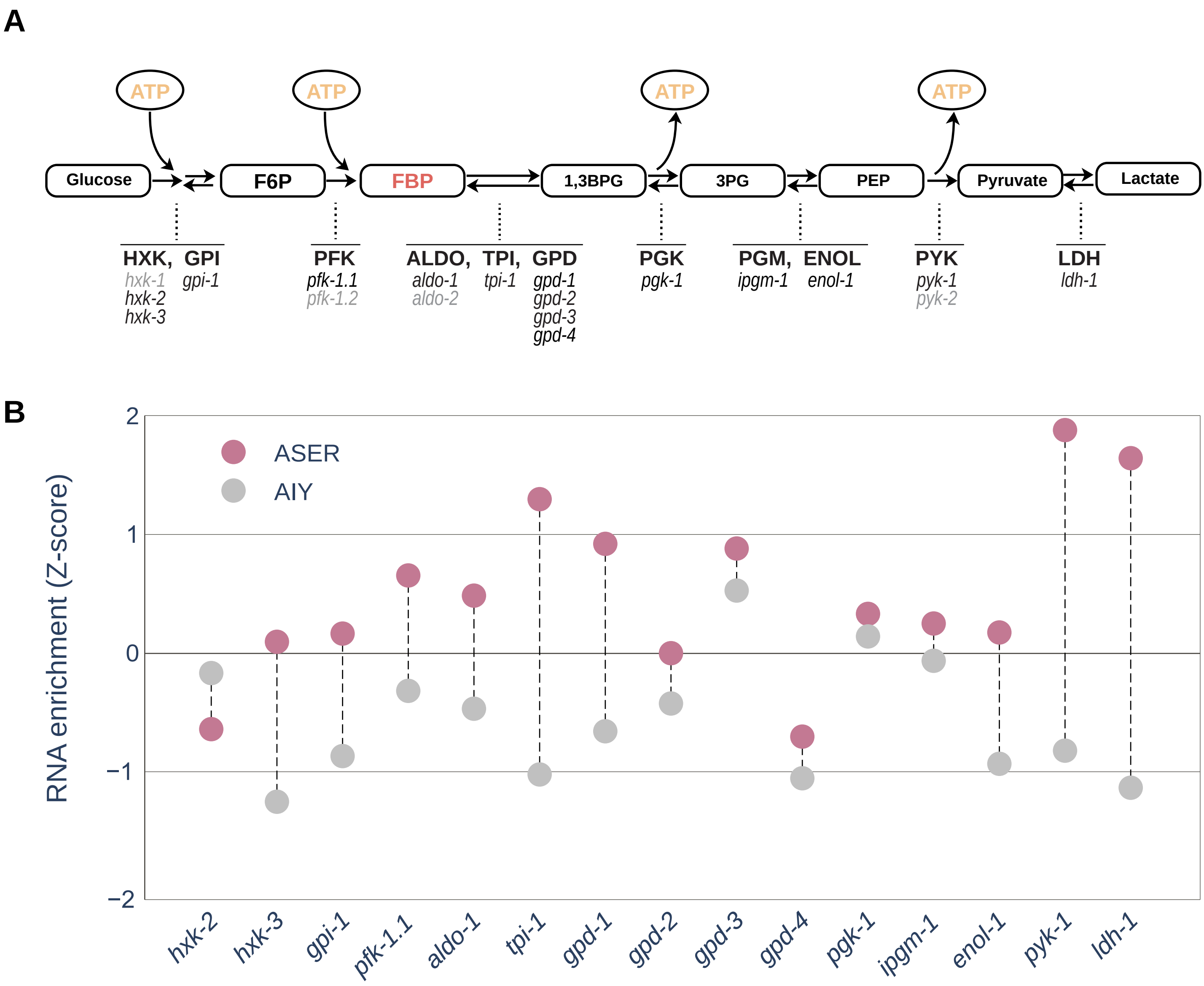
A
ATP
ATP
ATP
ATP
Glucose
F6P
FBP
1,3BPG
3PG
PEP
Pyruvate
Lactate
HXK, GPI
hxk-1 gpi-1
hxk-2
hxk-3
PFK
pfk-1.1
pfk-1.2
ALDO, TPI, GPD
aldo-1 tpi-1 gpd-1
aldo-2 gpd-2
gpd-3
gpd-4
PGK
pgk-1
PGM, ENOL
ipgm-1 enol-1
PYK
pyk-1
pyk-2
LDH
ldh-1
B
ASER
AIY
RNA enrichment (Z-score)
2
1
0
−1
−2
hxk-2
hxk-3
gpi-1
pfk-1.1
aldo-1
tpi-1
gpd-1
gpd-2
gpd-3
gpd-4
pgk-1
ipgm-1
enol-1
pyk-1
ldh-1

Figure 3-S2

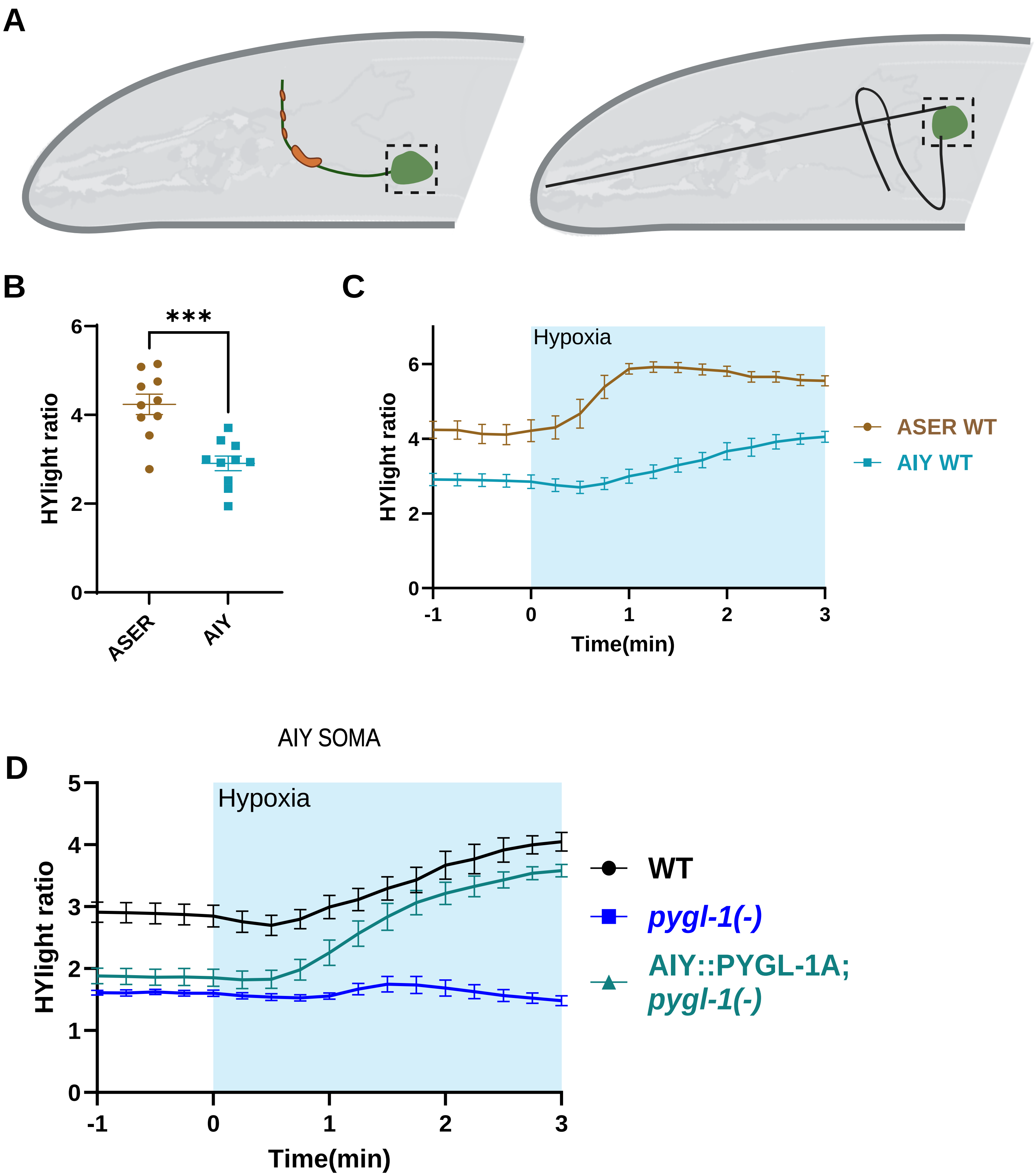
A
B
***
HYlight ratio
6
4
2
0
ASER
AIY
C
Hypoxia
HYlight ratio
6
4
2
0
-1
0
1
2
3
Time(min)
ASER WT
AIY WT
AIY SOMA
D
Hypoxia
HYlight ratio
5
4
3
2
1
0
-1
0
1
2
3
Time(min)
WT
pygl-1(-)
AIY::PYGL-1A;
pygl-1(-)

Figure 4-S1

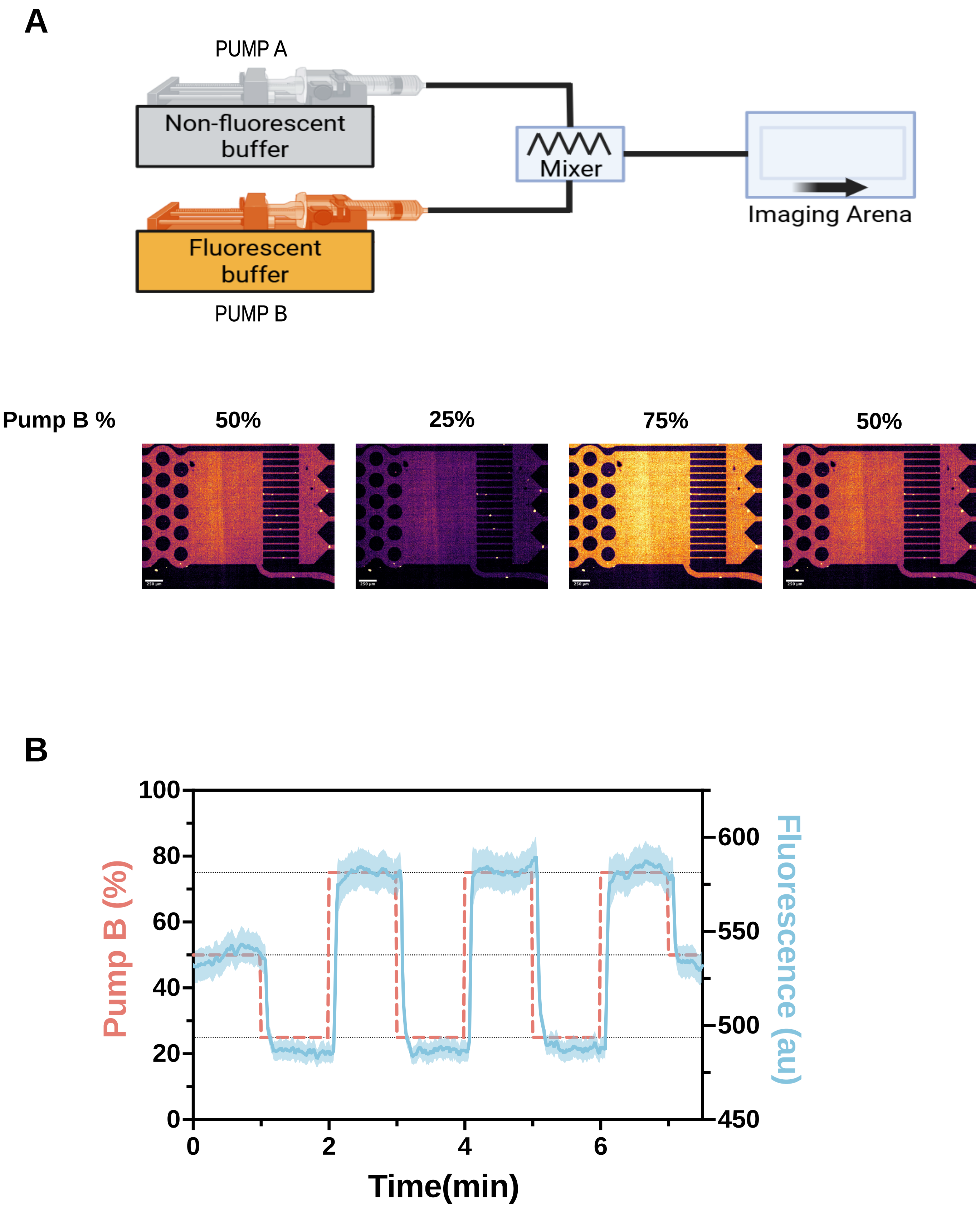

A
PUMP A
Non-fluorescent buffer
Mixer
Imaging Arena
Fluorescent buffer
PUMP B
Pump B %
50%
25%
75%
50%
B
Pump B (%)
Fluorescence (au)
Time(min)
0
20
40
60
80
100
450
500
550
600
0
2
4
6

Figure 4-S2

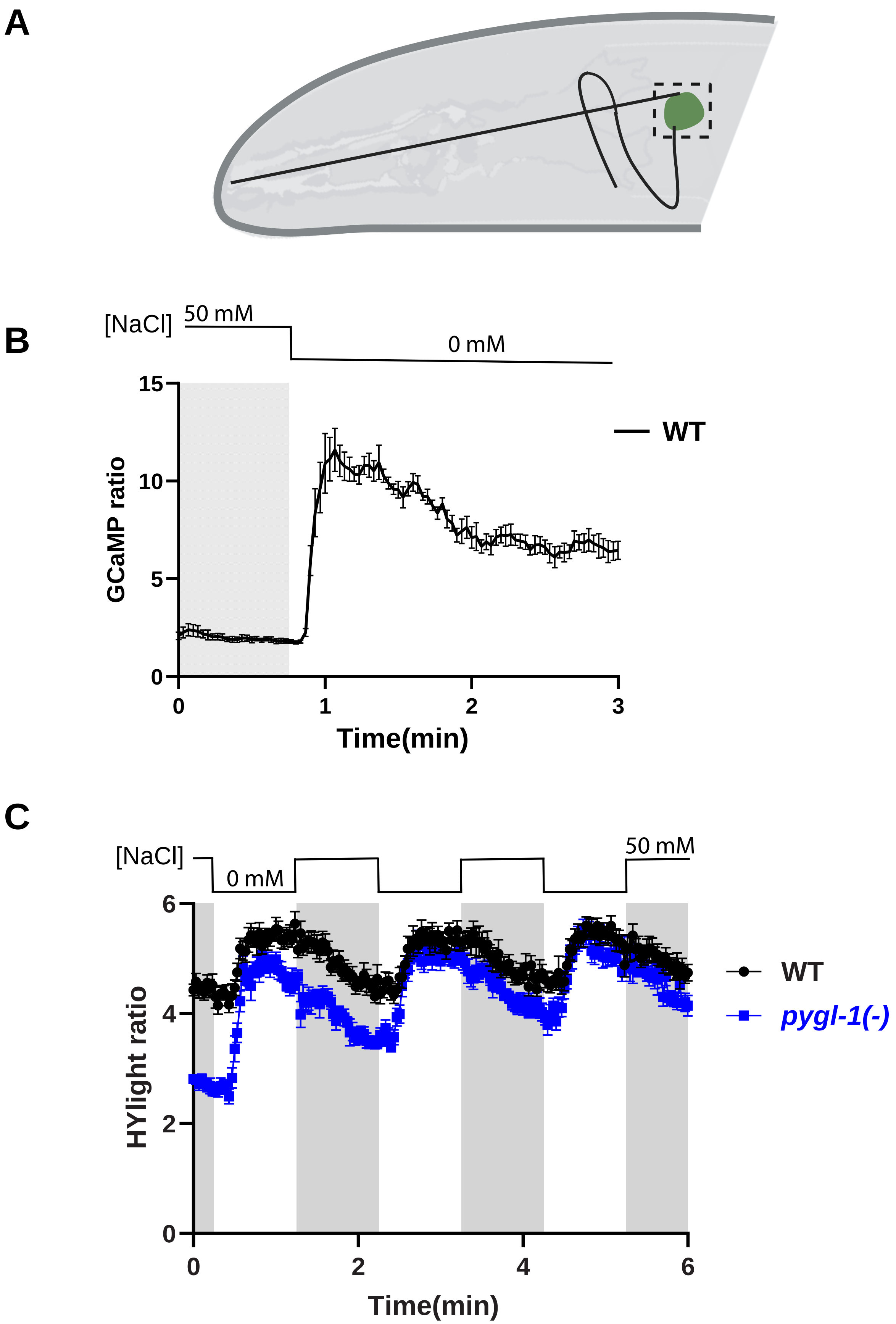
A
B
[NaCl]
50 mM
0 mM
15
10
5
0
GCaMP ratio
— WT
0
1
2
3
Time(min)
C
[NaCl]
0 mM
50 mM
6
4
2
0
HYlight ratio
WT
pygl-1(-)
0
2
4
6
Time(min)
